# Deep learning classification of canine behavior using a single collar-mounted accelerometer: Real-world validation

**DOI:** 10.1101/2020.12.14.422660

**Authors:** Robert D. Chambers, Nathanael C. Yoder, Aletha B. Carson, Christian Junge, David E. Allen, Laura M. Prescott, Sophie Bradley, Garrett Wymore, Kevin Lloyd, Scott Lyle

**Affiliations:** Pet Insight Project, Kinship, San Francisco, California, United States of America; WALTHAM Petcare Science Institute, Melton Mowbray, Leicestershire, United Kingdom

## Abstract

Collar-mounted canine activity monitors can use accelerometer data to estimate dog activity levels, step counts, and distance traveled. With recent advances in machine learning and embedded computing, much more nuanced and accurate behavior classification has become possible, giving these affordable consumer devices the potential to improve the efficiency and effectiveness of pet healthcare. Here we describe a novel deep learning algorithm that classifies dog behavior at sub-second resolution using commercial pet activity monitors. We built machine learning training databases from over 5,000 videos of over 2,500 dogs and ran the algorithms in production on over 11 million days of device data. We then surveyed project participants representing 10,550 dogs, they provided us 163,110 event responses to validate real-world detection of eating and drinking behavior. The resultant algorithm displayed a sensitivity and specificity for detecting drinking behavior (0.949 and 0.999, respectively) and eating behavior (0.988, 0.983). We also demonstrated detection of licking, petting, rubbing, scratching, and sniffing. We show that the devices’ position on the collar has no measurable impact on performance. In production, users reported a true positive rate of 95.3% for eating (among 1,514 users), and of 94.9% (among 1,491 users) for drinking. The study demonstrates the accurate detection of important health-related canine behaviors using a collar-mounted accelerometer. We trained and validated our algorithms on a large and realistic training dataset, and we assessed and confirmed accuracy in production via user validation.

## Introduction

Much as recent progress in smartwatches has enabled new telehealth applications [1–4], recent progress in internet-connected pet wearables, such as collar-mounted activity monitors, has prompted interest in using these devices to improve the cost and efficacy of veterinary care [5]. Just as with smartwatches in human telehealth, accelerometer-based activity monitors have emerged as an inexpensive, low-power, and information-rich approach to pet health monitoring [6–8].

Accelerometer-based pet activity monitors analyze the moment-to-moment movement measured by a battery-powered accelerometer. They are typically attached to the pet via a collar, though attachment methods may be more elaborate in research settings. Using the device’s accelerometer signal (sometimes in combination with gyroscope, magnetometer, GPS, or other sensor signals), collar-mounted activity monitors can accurately estimate pet activity levels [9–14], step count, and distance traveled [12].

In recent years, advances in machine learning have allowed pet activity monitors to move beyond estimating aggregate activity amounts, to detecting when and for how long a pet performs common activities such as walking, running, lying down, or resting [15–17], These biometric capabilities have progressed to include increasingly specific and varied activities such as drinking, eating, scratching, and head-shaking [8,16,18–21].

The benefits of accurate and quantitative behavior detection in pet health are extensive. Pet activity monitors have been shown to be useful in the detection and diagnosis of pruritis [22,23] and in potential early prediction of obesity [24]. They have also been used in monitoring response to treatments such as chemotherapy [25]. Furthermore, statistical analysis of activity and behavior monitoring on large numbers of pets can be an expedient approach to medical and demographic studies [24], since sample sizes are potentially very large.

Although several studies have demonstrated and measured the accuracy of activity recognition algorithms [8,16,18–21], the datasets used to train and evaluate the algorithms are typically not representative of the broad range of challenging environments in which commercial pet activity monitors must function. For instance, most existing studies use exclusively healthy dogs and are often run in controlled environments that promote well-defined and easily detectable behaviors with a low risk of confounding activities.

Unfortunately, real-world algorithm performance often lags far behind the performance measured in controlled environments [26,27]. For instance, existing studies typically ensure careful installation of the device in a specific position on a properly adjusted collar. In real-world usage, collars vary in tightness and often rotate to arbitrary positions unless the activity monitor device is very heavy. Collar rotation and tightness [28], as well as the use of collar-attached leashes [29], can compromise performance. In our experience, confounding activities like riding in a car or playing with other pets can produce anomalous results if not adequately represented in training datasets. Finally, some studies use multiple accelerometers or harness-mounted devices [30], which limit applicability in many consumer settings.

The work described here was performed as part of the Pet Insight (PI) Project [31], a large pet health study to enable commercial pet activity monitors to better measure and predict changes in a pet’s health by:

1. Sourcing training data from project participants and external collaborators to build machine learning training databases and behavior detection models such as those described in this work.
2. Combining activity data, electronic medical records, and feedback from more than 69,000 devices distributed to participants over 2-3 years to develop and validate proactive health tools.
3. Using the resulting datasets, currently covering over 11 million days in dogs’ lives, to enable insights that support pet wellness and improve veterinary care.

This work presents the results of the PI Project’s efforts to develop and validate these behavior classification models [32]. It includes evaluation of model performance in a real-world context and addresses limitations from controlled research settings such as device fit and orientation.

## Materials and methods

### Activity Monitor

Data were collected primarily via a lightweight canine activity monitor (Whistle FIT^®^, Mars Petcare, McLean, VA, Fig 1), which was designed and produced specifically for this study). Smaller amounts of data were collected via the commercially available Whistle 3^®^ and Whistle GO^®^ canine activity monitors. All three devices used the same accelerometer. Unlike the Whistle FIT^®^, these latter devices are furnished with GPS receivers and cellular radios. However, in all cases, the behavior classification in this study is performed using only the output of the devices’ 3-axis accelerometers.

**Fig 1.**
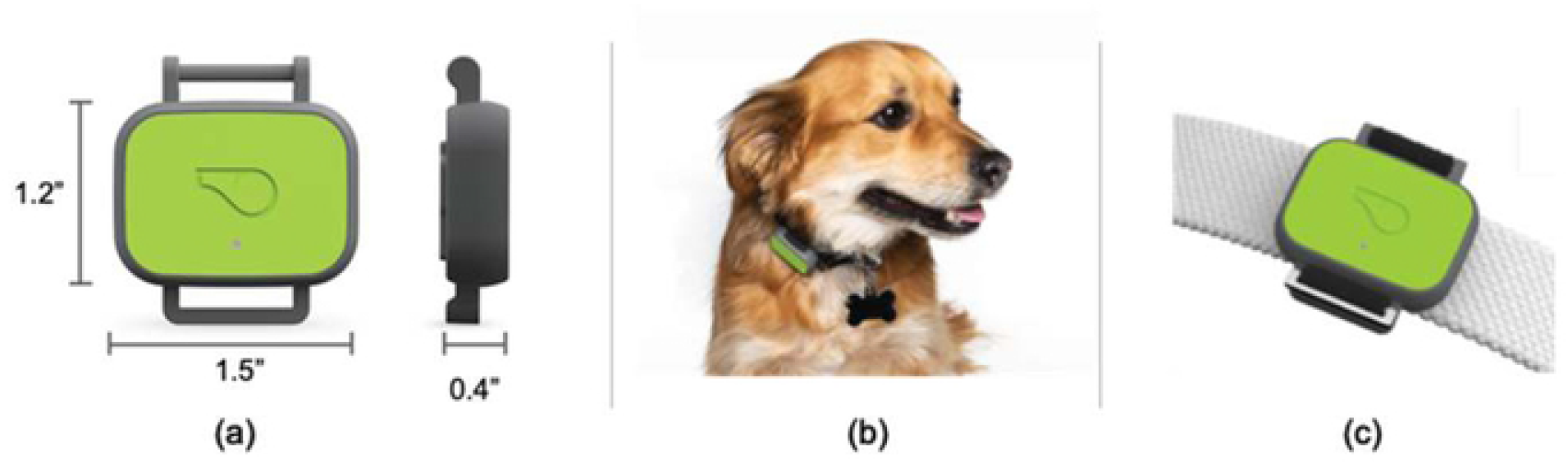
Activity monitors used in this study. Most data in this study were acquired from Whistle FIT^®^ activity monitors. Device dimensions are shown in (a), and a device in use is shown in (b). The device often rotates to different positions around each dog’s collar. The device can attach to most dog collars up to 1” (25 mm). Attachment detail is shown in (c). The two other devices used this study (the Whistle 3^®^ and the Whistle GO^®^) are larger and heavier.

### Accelerometry data collection

All monitoring devices acquired accelerometry data and uploaded it according to their usual operation. That is, the devices acquired 25-50 Hz 3-axis accelerometry data for at least several seconds whenever significant movement was detected. Data were compressed and annotated with timing data using a proprietary algorithm. Data were temporarily stored on-device and then uploaded at regular intervals when the devices were in Wi-Fi range. Uploads were processed, cataloged, and stored in cloud-hosted database services by Whistle servers. The compressed accelerometry data were retrieved on demand from the cloud database services in order to create the training, validation, and testing databases used in this study.

### Animal behavior data collection

Animal behavior data was collected (summarized in Table 1 and described further elsewhere in this report) and used to create two datasets used in model training and evaluation:

1. **Crowd-sourced** (*crowd*) **dataset**. This dataset contained both (a) long (multi-hour) in-clinic recordings, as well as (b) shorter recordings submitted by project participants. This large and diverse dataset was meant to reflect real-world usage as accurately as possible.
2. **Eating and drinking** (*eat/drink*) **dataset**. This dataset consisted of research grade sensor and data using a protocol designed to represent EAT and DRINK behaviors. Other observed behaviors were incidental.

**Table 2.** Datasets Derived from Animal Data

| Dataset | Protocol<br>Name | Location | Description |
| --- | --- | --- | --- |
| Crowd | In-Clinic<br>Crowd-Sourcing | Banfield Pet<br>Hospital<br>clinics | Dogs that were awaiting in-clinic care were outfitted with activity monitors and were video recorded for several hours each performing typical in-kennel behaviors. |
|  | PI<br>Participant<br>Crowd-Sourcing | Multi-source:<br>For instance,<br>at-home, in-<br>car, during<br>walks and<br>hikes. | Participants in the Pet Insight<br>Project used smartphones to<br>video record their dogs while<br>wearing activity monitors in<br>every-day situations. Collar fit,<br>device orientation, environment,<br>and animal behavior were<br>meant to be representative of<br>real-world usage. |
| Eat/drink | Waltham<br>Eat/Drink<br>Study | WALTHAM<br>Petcare<br>Science<br>Institute | Dogs were video recorded<br>eating and drinking by<br>researchers at the WALTHAM<br>Petcare Science Institute.<br>Collar fit and orientation were<br>controlled, and dog behaviors<br>exhibited were relatively<br>consistent. |

For brevity, we refer to these datasets simply as the *crowd* and *eat/drink* datasets.

### Eat/drink study protocol

This study was conducted using dogs owned by the WALTHAM Petcare Science Institute and housed in accordance with conditions stipulated under the UK Animals (Scientific Procedures) Act 1986. Briefly, the dogs were pair housed in environmentally enriched kennels designed to provide dogs free access to a temperature-controlled interior and an external pen at ambient temperature. Dogs were provided with sleeping platforms at night. The dogs had access to environmentally enriched paddocks for group socialization and received lead walks and off-lead exercise opportunities during the day. Water was freely available at all times and dogs were fed to maintain an ideal body condition score. The study was approved by the WALTHAM Animal Welfare and Ethical Review Body. One hundred and thirty-eight dogs across 5 different breeds (72 Labrador Retrievers, 18 Beagles, 17 Petit Basset Griffon Vendeens, 14 Norfolk Terriers and 17 Yorkshire Terriers) took part for two consecutive days each. Each dog was recorded once a day during its normal eating and drinking routine using a GoPro camera (GoPro, San Mateo, CA).

In this study, either one (ventral only) or four (ventral, dorsal, left, and right) activity monitors were affixed to a collar. For each observation, the collar was removed from the dog, the correct number of activity monitors were attached, and then shaken sharply in view of the camera to provide a synchronization point that was identifiable in both the video and accelerometer signals (so that any time offset could be removed). The collar was then placed on the dog at a standardized tightness. The dogs were recorded from approximately one minute before feeding until approximately one minute after feeding. In order to increase the diversity of the dataset, collar tightness was varied between a two-finger gap and a four-finger gap, and food bowls were rotated between normal bowls and slow-feeder or puzzle-feeder bowls. For each data recording, researchers noted the date and time, device serial number(s), collar tightness, food amount and type, and various dog demographic data.

### Crowd-sourcing protocol

Pet Insight participants were requested to use smartphones to video record their pets performing everyday activities while wearing activity monitors. The participants were told that the activity monitor should be worn on the collar but were not given any other instructions about how the collar or monitor should be worn. Participants were asked to prioritize recording health-related behaviors like scratching or vomiting, but to never induce these events and to never delay treatment in order to record the behaviors. As a participation incentive, for every crowd-sourced video used, the PI project donated one dollar to a pet-related charity.

After recording each video, participants logged into the PI crowd-sourcing website, provided informed consent, uploaded the recorded video, and completed a short questionnaire confirming which pet was recorded and whether certain behaviors were observed. The device automatically uploaded its accelerometry data to Whistle servers.

### In-clinic observational protocol

This study was conducted at several Banfield Pet Hospital (BPH) clinics. Its objective was to acquire long-duration (multi-hour) naturalistic recordings to augment the shorter crowd-sourced recordings, which were typically several minutes or less in duration.

Randomly selected BPH clients who chose to participate signed fully informed consent forms. Their dogs were outfitted with Velcro breakaway collars with one attached activity monitor device each. Collar tightness and orientation were not carefully controlled. Video was recorded via a 4-channel closed-circuit 720p digital video security system. Video cameras were ceiling- or wall-mounted and oriented towards the in-clinic kennels so that up to four dogs could be observed at a time. For each recording, researchers noted the date and time, the device serial number, and the dog/patient ID number.

### Video labeling

All uploaded videos were transcoded into a common format (H.264-encoded, 720p resolution, and up to 1.6 Mb/s) using Amazon’s managed Elastic Transcoder service, and their audio was stripped for privacy. Video start times were extracted from the video metadata and video filenames. Matching device accelerometry data was downloaded from Whistle’s databases, and automatic quality checks were performed.

Videos were then labeled by trained contractors using the open-source BORIS (Behavioral Observation Research Initiative Software) software application [33]. The resulting event labels were imported and quality-checked using custom Python scripts running on one of the PI project’s cloud-based web servers. Labels were stored alongside video and participant metadata in a PostgreSQL database.

All video labeling contractors were trained using a standardized training protocol, and inter-rater reliability analyses were performed during training to ensure consistent labeling. Videos were labeled according to a project ethogram [8,15,20]. This report describes several of these label categories.

Labelers divided each video into *valid* and *invalid* regions. Regions were only valid if the dog was clearly wearing an activity monitor and was fully and clearly visible in the video. Invalid regions were subsequently ignored. In each *valid* video region, the labeler recorded exactly one *posture,* and any number (0 or more) of applicable *behaviors.*

*Postures* (Table 2) reflect the approximate position and energy expenditure level of the pet, while *behaviors* (Table 3) characterize the pet’s dominant behavior or activity in a given moment. For instance, during a meal, a dog might exhibit a STAND posture and an EAT behavior. While pausing afterwards, the same dog might exhibit a STAND posture and no behavior. Multiple simultaneous behaviors are rare but possible, such as simultaneous SCRATCH and SHAKE behaviors.

**Table 2.** Postures ethogram.

| Posture | Activity |
| --- | --- |
| LIE DOWN | Lying down. |
| SIT | Sitting still with little to no movement |
| STAND | Standing still with little to no movement. |
| WALK | Purposeful walking from one point to another. |
| VIGOROUS | Catch-all for high-energy activities such as running, swimming, and playing. |
| MIXED | Default category for any other posture, for ambiguous postures, and for postures that are difficult to label due to rapid changes. |

**Table 3.**
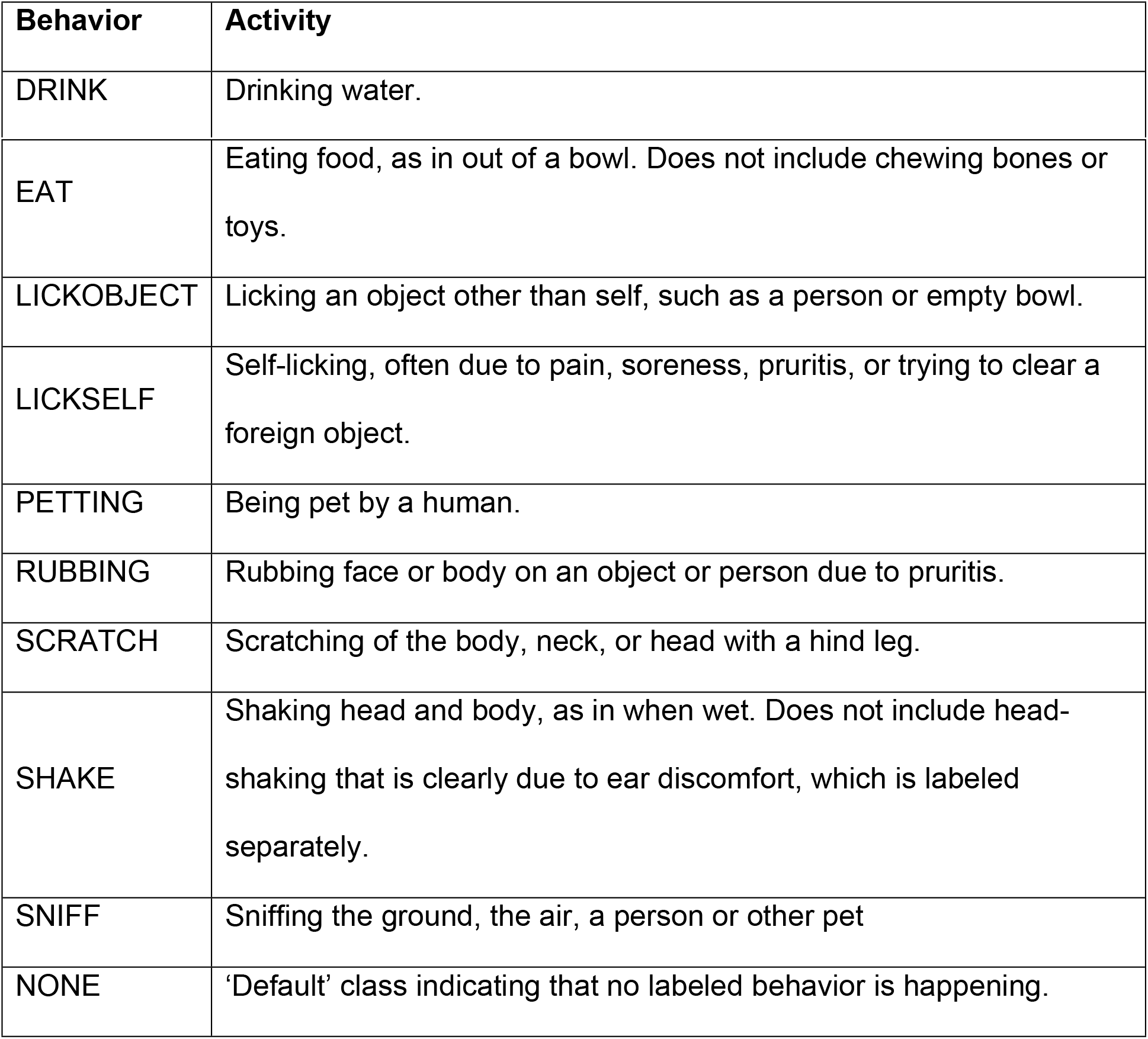
Behaviors ethogram.

### Training data preparation

Although accelerometer data and smartphone video data were both time-stamped using the devices’ network-connected clocks, inaccuracies led to alignment errors of typically several seconds, and sometimes much longer. Short activities such as SHAKE, in particular, require more accurate alignment. We aligned approximately 1,200 videos manually by matching peaks in accelerometer activity to labels for high-intensity behaviors like SHAKE and SCRATCH. We used these manual alignments to develop and validate an automatic alignment algorithm that aligned the remaining videos.

We created each of the two training datasets *(crowd* and *eat/drink)* by:

1. Selecting appropriate videos from our database.
2. Limiting the number of entries per dog to 30 (some dogs are overrepresented in our database).
3. Allocating all of each dog’s data into one of 5 disjoint cross-validation folds.
4. Downloading each dataset and labeling each time-point with a posture and/or behavior(s).

The specific method of separating data into cross-validation folds (step 3 above) is critical [34]. Classifiers trained on individual dogs have been shown to overperform on those dogs relative to others, even if those classifiers are trained and evaluated using separate experimental observations. Gerencsér et. al. experienced an accuracy reduction from 91% for a single-subject classifier to 70-74% when generalizing to other dogs [35]. Consequently, we were careful to ensure that all of a dog’s videos fall in a single fold, so that data from a single dog is never used to both train and evaluate a classifier.

The overall data acquisition process, from video capture to a completed dataset, is shown in Fig 2.

**Fig 2.**
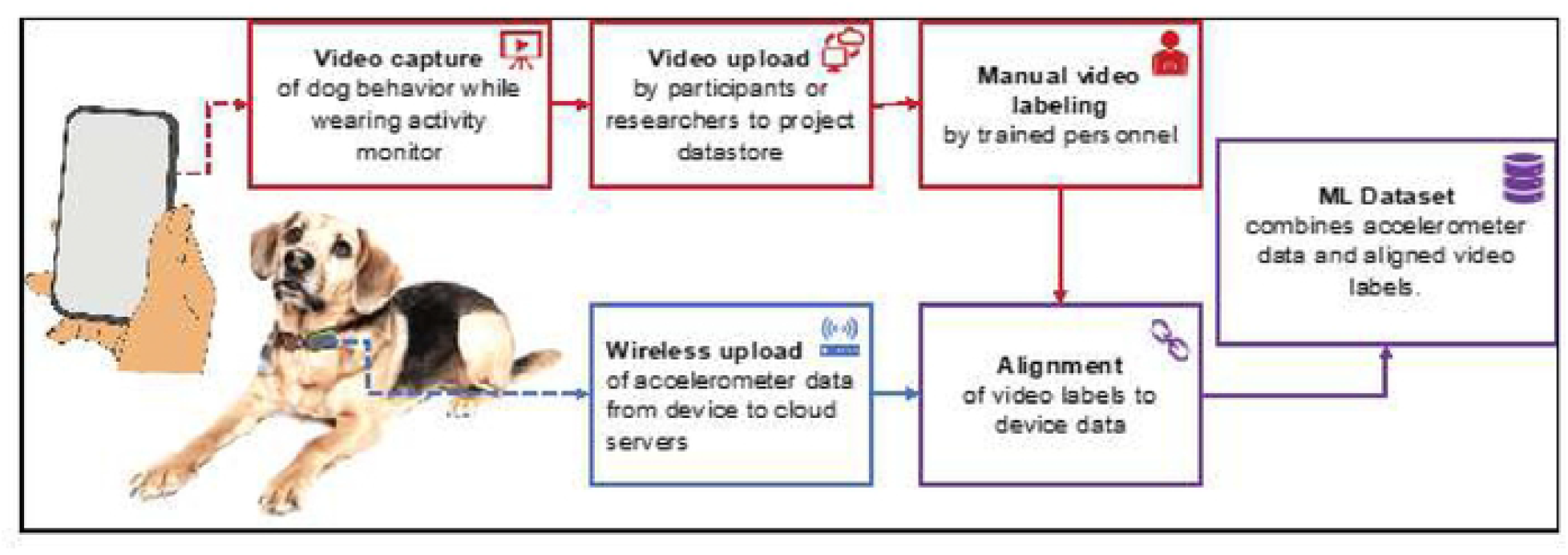
Data acquisition flow. Dogs wearing collar-mounted activity monitors were video recorded performing behaviors of interest or performing everyday activities. Videos were uploaded and the behaviors exhibited in them were manually labeled (tagged). The devices automatically uploaded accelerometer (activity) data to cloud servers, and the device data was aligned with the video labels to remove any temporal offset. The aligned labels and accelerometer time series were combined into datasets suitable for training machine learning models.

### Deep learning classifier

Our deep learning classifier is based on our FilterNet architecture, which we have published in detail in a previous work [32]. We implemented the model in Python using PyTorch v1.0.1 [36] and the 2020.02 release of the Anaconda Python distribution (64-bit, Python 3.7.5). We trained and evaluated our models on p2.xlarge instances on Amazon Web Services [37] with 4 vCPUs (Intel Xeon E5-2686 v4), 61 GB RAM, and a NVIDIA Tesla k80 GPU with 12 Gb RAM, running Ubuntu 18.04.4.

We used the *crowd* dataset for cross-validated training and evaluation (Fig 3). Specifically, we trained and evaluated five different models, using a different held-out fold as a test set for each model. We combine the models’ predictions for each of the five test sets for model evaluation, as described below. We also generated behavior classifications for the *eat/drink* dataset using one of the models trained on the *crowd* dataset (that is, we did not use the *eat/drink* dataset for model training). There were no dogs in common between the *crowd* and *eat/drink* datasets, so cross-validation was not needed in this step.

**Fig 3.**
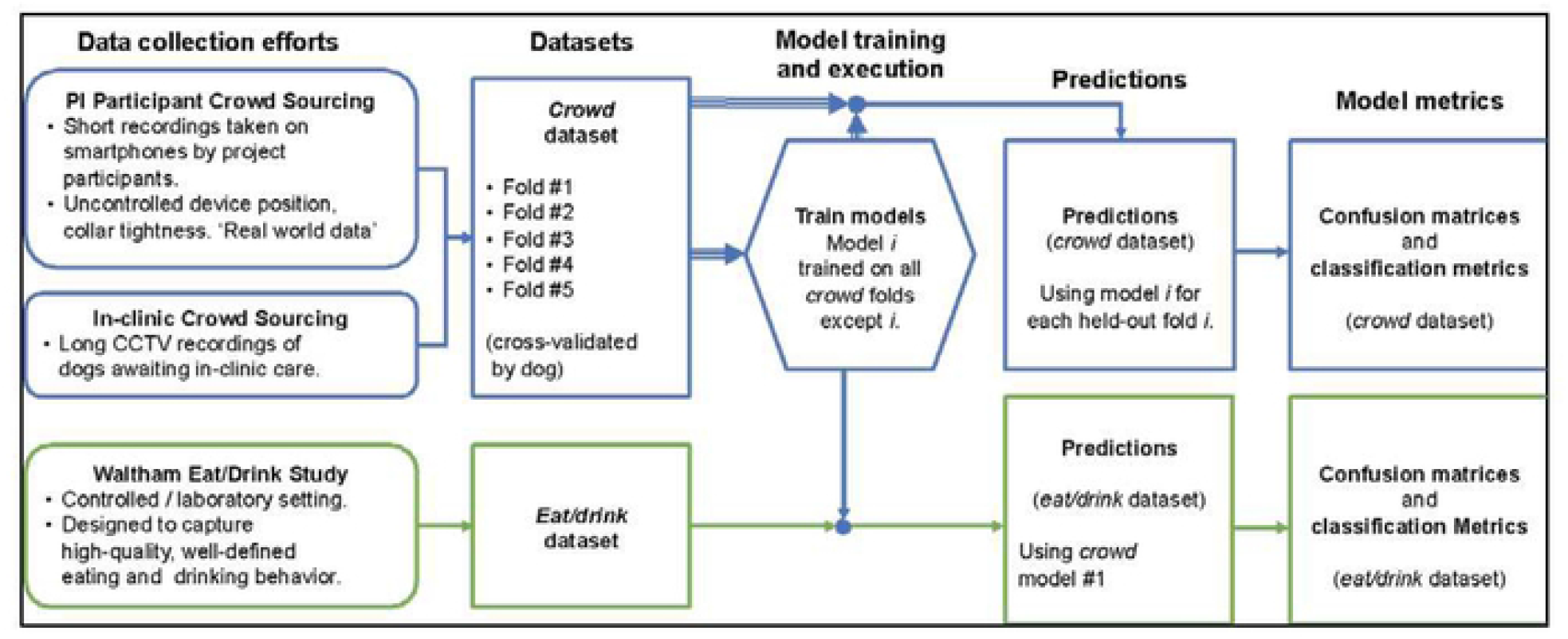
Model training and evaluation data flow. The *crowd* dataset consisted of naturalistic, highly diverse data divided by dog into five folds. The *eat/drink* dataset focused on high-quality eating and drinking data. Behavior classification models were trained and evaluated in a cross-validated fashion on the *crowd* dataset, and the first of these five models was also evaluated on the *eat/drink* dataset. Confusion matrices and classification metrics were produced for each dataset using the resulting predictions.

### Evaluation

For evaluation, we modeled the task as two multi-class classification problems, one for behaviors and one for postures. At each timepoint in each video entry in a dataset we recorded the *labeled* behavior and posture, and every 320 ms we calculate the most likely *predicted* behavior and posture. We tallied the labeled and predicted pairs from all five test folds together using the *PyCM* multiclass confusion matrix library to create separate behavior and posture confusion matrices [38]. We used the *PyCM* package to calculate metrics derived from the confusion matrices [39].

Because the *MIXED* posture is used primarily for expediency in labeling, we dropped any timepoints with *MIXED* labels from the postures confusion matrix, and replaced any MIXED-class posture predictions with the next most likely prediction for that timepoint. We also excluded any timepoints with more than one simultaneous labeled behavior (about 3% of the data) from the behaviors confusion matrix.

Furthermore, following Uijl et. al. [8], we excluded any timepoints within 1 second of a class transition in both classification problems. However, also similar to [8], we treated the SHAKE class differently due to its very short duration. For SHAKE, we only excluded the outer one-third second. In dropping these transition regions, we attemptted to follow established convention for minimizing the effects of misalignment in labeling, and to make our reported results easier to compare to related works.

### User validation

Although the *crowd* dataset is meant to be representative of real-world data, it is subject to biases such as underrepresentation of behaviors that are unlikely to be video recorded, such as riding in cars or staying at home alone. Furthermore, it is impossible to anticipate all of the myriad situations that may serve as confounders.

Consequently, we ran real-world user validation campaigns on the two behaviors that users are most likely to be aware of, EAT behavior and DRINK behavior. We defined events as periods of relatively sustained, specific behaviors detected with high confidence, such as eating events (meals) consisting of several minutes of sustained eating behavior. We adapted our production system, which runs the models described in this work in nearreal-time on all PI project participants, to occasionally send validation emails to participants when an EAT or DRINK event had occurred within the past 15 minutes. Respondents categorized the event detection as correct (“Yes”) or incorrect (“No”) or indicated that they weren’t sure. Users were able to suggest what confounding event may have triggered any false predictions. We excluded any responses that arrived more than 60 minutes after an event’s end, as well as any “Not Sure” responses.

## Results

### Data collected

After applying the steps described above the *crowd* dataset contained data from 5,063 videos representing 2,217 subjects, and the *eat/drink* dataset contained data from 262 videos representing 149 unique dogs.The distribution of weights and ages represented in these datasets is shown in Fig 4, while a breed breakdown is given in Table 4. As expected, the crowd dataset exhibits a far greater diversity of weights, ages, and breeds than the eat/drink dataset, since the eat/drink subjects are sampled from several relatively homogeneous subpopulations.

**Fig 4.**
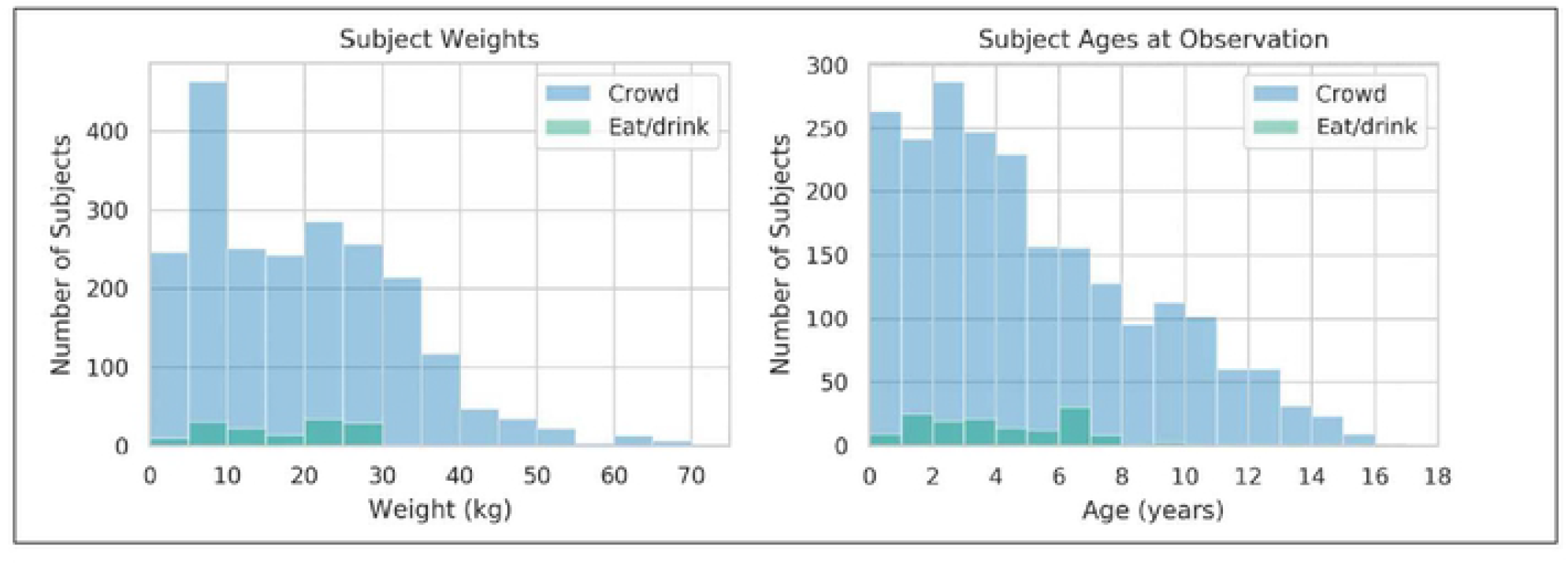
Weight and age distribution for the *crowd* and *eat/drink* datasets. Due to its more heterogenous population, the crowd dataset had much greater variability than the eat/drink dataset.

**Table 4.**
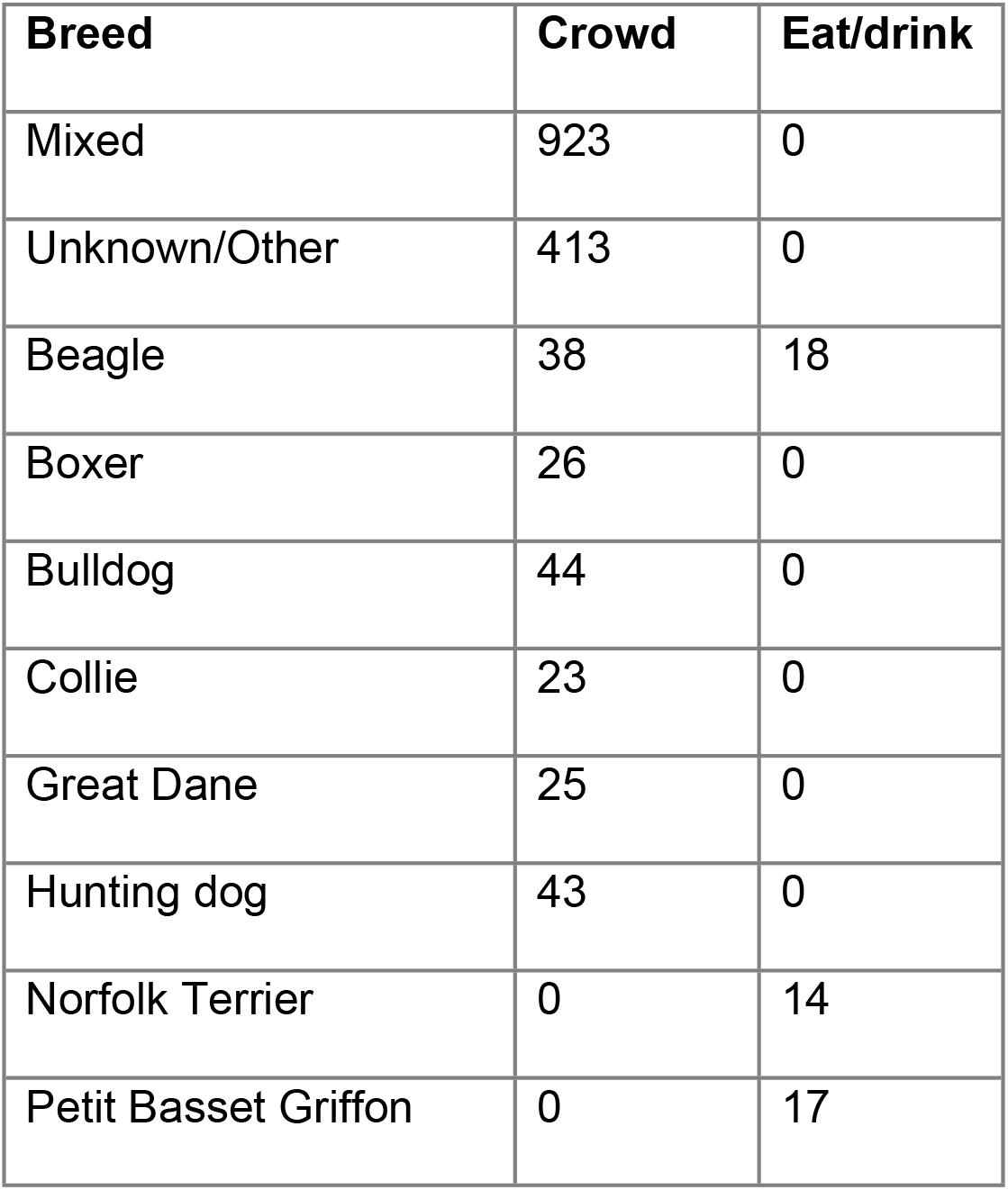
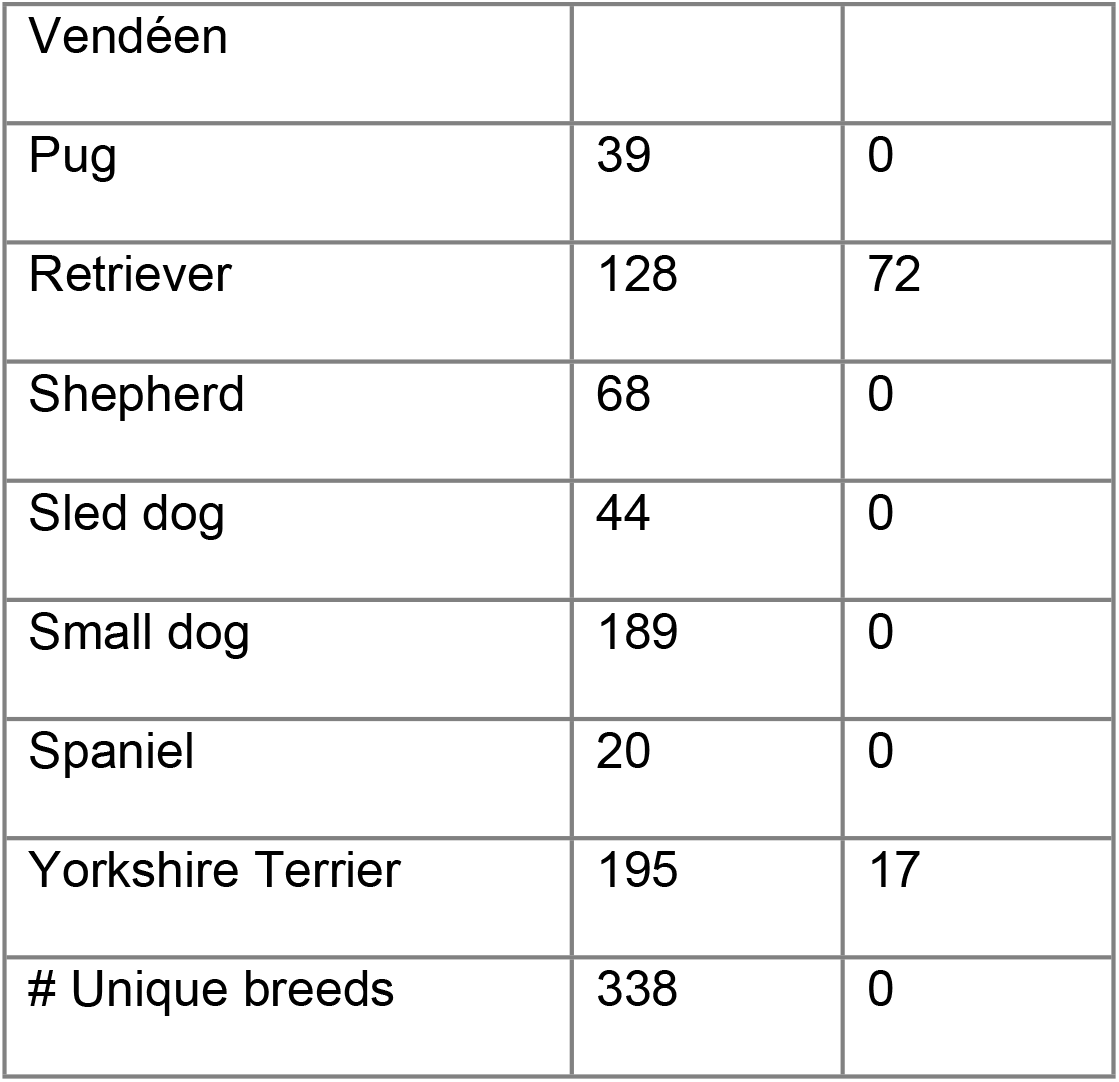
Breed or breed category breakdown for the *crowd* and *eat/drink* datasets.

These datasets also differed in the length and frequency of labeled events, as shown in Table 5. The *crowd* and *eat/drink* datasets contain 163.9 and 22.4 hours of video data labeled as VALID, respectively.

**Table 5.**
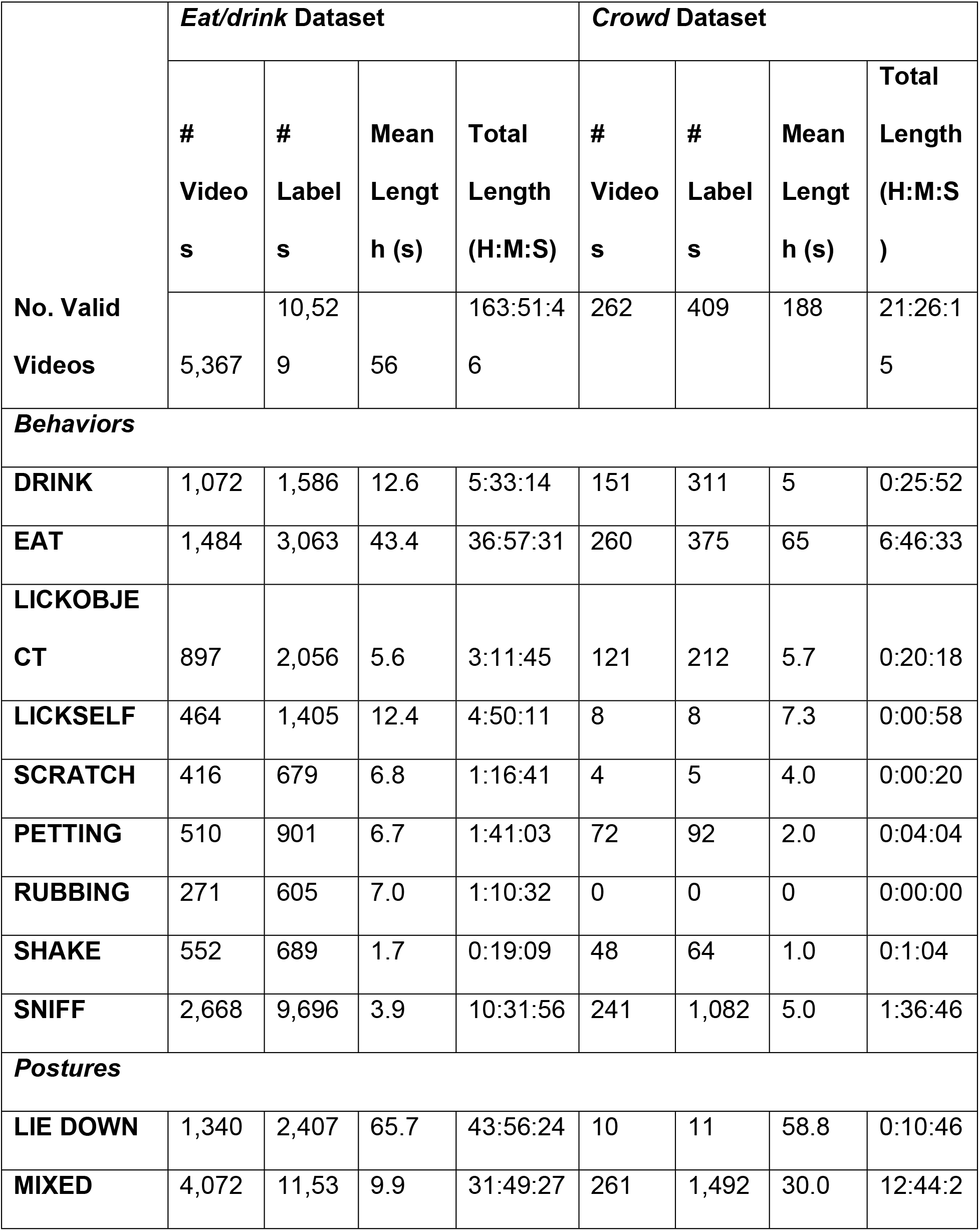
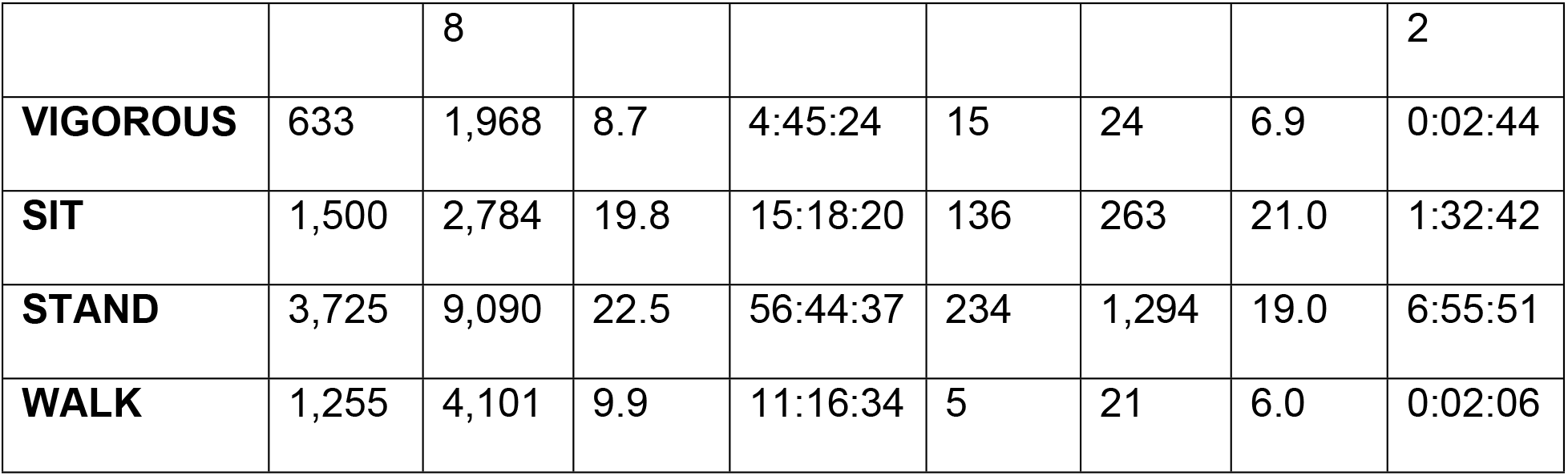
Summary of labeled *crowd* and *eat/drink* datasets.

The EAT class was highly represented in both the *crowd* dataset (because participants were specifically requested to submit videos of their dogs at mealtime, since it is an easily filmed and important behavior) and in the *eat/drink* dataset (due to study design). The *eat/drink* dataset included only small amounts of incidental LICKSELF, SCRATCH, PETTING, and SHAKE behavior, while the *crowd* dataset contained many of these events because participants were repeatedly reminded of their importance.

The class distribution of both datasets is highly imbalanced, which presented a challenge for algorithm training. For instance, in the *crowd* dataset, which we used for training, the EAT class total duration is 117 times greater than that of SHAKE.

The distribution of lengths for each label class was highly skewed, with many short labels and a smaller number of longer labels (Fig 5). Some of this skew was due to label fragmentation, where a long stretch of the labeled activity is interrupted either by the dog temporarily pausing (for instance, lifting up its head to look around several times while drinking or while eating a meal) or by discontinuities in the labeling when the dog leaves the camera’s field of view (since labelers only marked videos as VALID when the dog was fully and clearly visible). The distribution of SHAKE labels was less skewed, likely because it is typically a short behavior and less prone to interruption.

**Fig 5.**
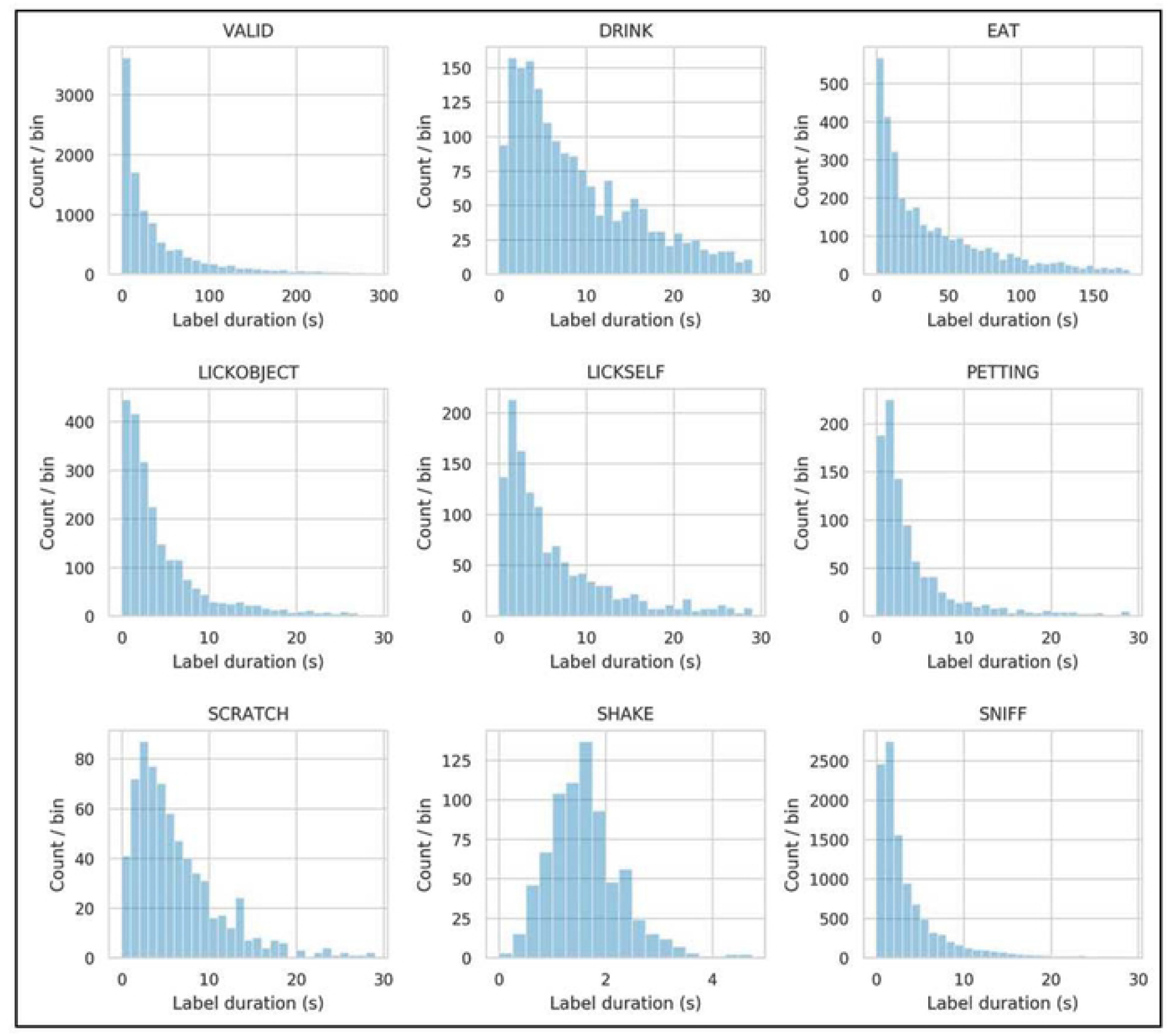
Distribution of label durations in the combined *eat/drink* and *crowd* datasets. Except for SHAKE, all labels exhibit a highly skewed distribution, probably the result of some longer label segments becoming fragmented.

### Classification accuracy

Cross-validated classification metrics for the *crowd* dataset are given in Table 6, and classification metrics obtained from evaluating the *eat/drink* dataset using a model trained on the *crowd* dataset are given in Table 7. The tables give metrics both for behavior and posture classes. However, some subsequent sections report only behaviors, because postures are much less carefully labeled and are typically used in an aggregate form where individual misclassifications are less important.

It is important to note that the class balance (class prevalence) of these datasets is not representative of real-world canine behavior. Because the videos are typically taken in stimulating or interesting situations, these datasets exhibit a lower relative prevalence of LIE DOWN and other low-energy postures. Furthermore, the datasets exhibit much higher levels of EAT, DRINK, and possibly other behaviors, due to either study design (in the *eat/drink* dataset) or because the PI project requested that participants film certain behaviors.

Of the metrics in Table 6 and Table 7, only sensitivity and specificity are independent of class prevalence.

**Table 6.**
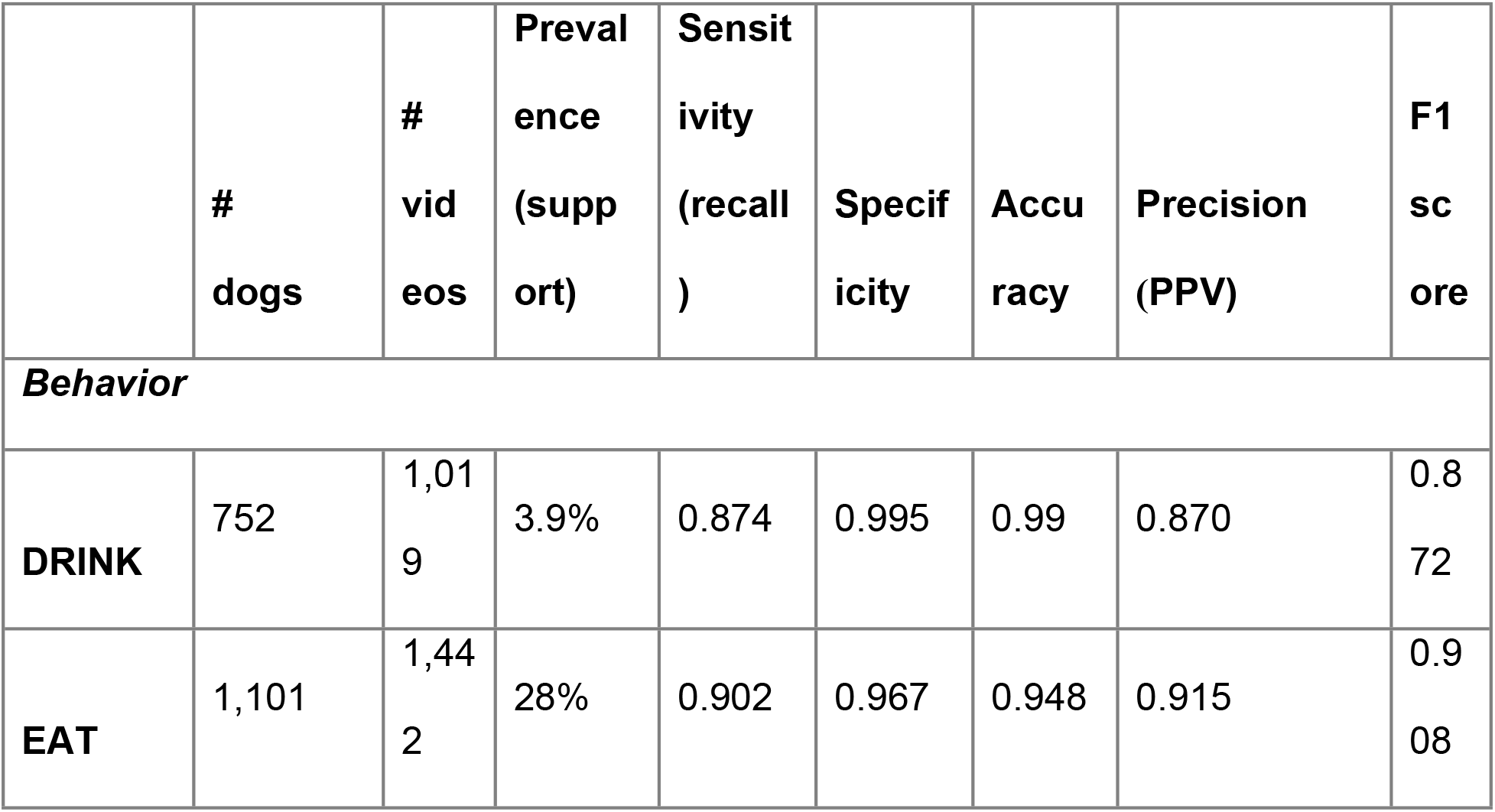
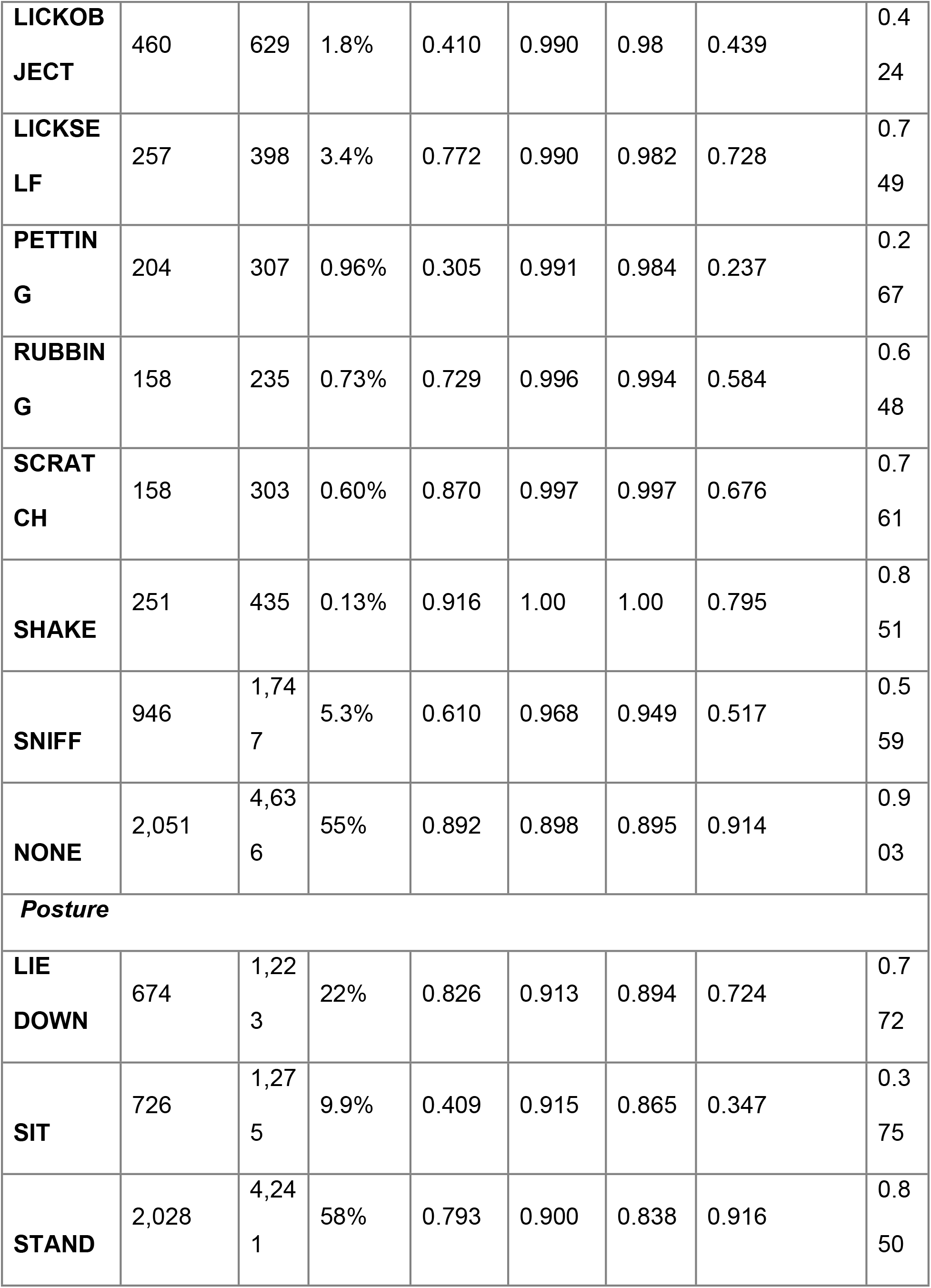
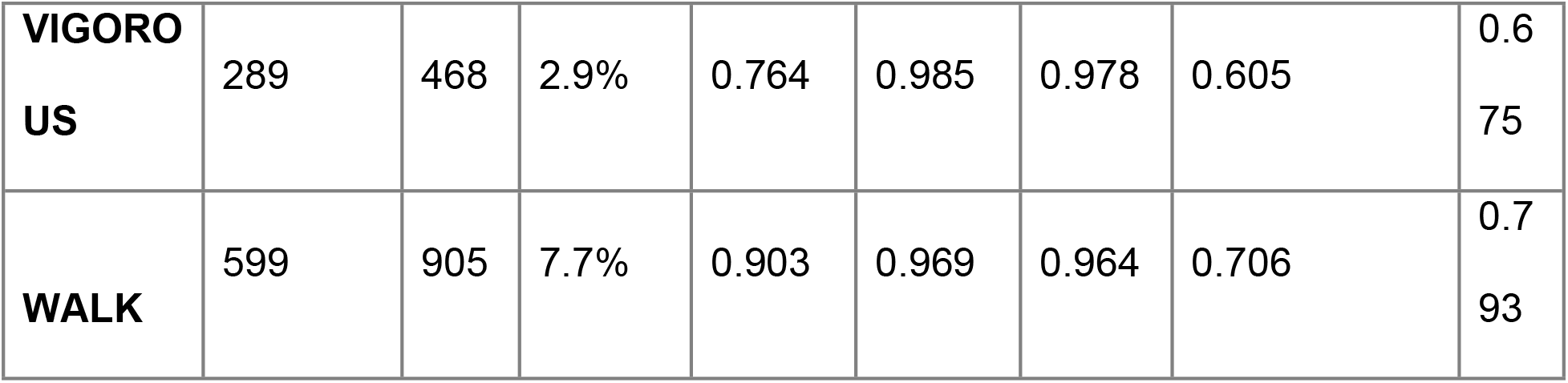
Classification metrics for the *crowd* dataset.

|  | #<br>dogs | #<br>videos | Prevalence<br>(support) | Sensitivity<br>(recall) | Specificity | Accuracy | Precision<br>(PPV) | F1<br>score |
| --- | --- | --- | --- | --- | --- | --- | --- | --- |
| ***Behavior*** |  |  |  |  |  |  |  |  |
| **DRINK** | 752 | 1,019 | 3.9% | 0.874 | 0.995 | 0.99 | 0.870 | 0.872 |
| **EAT** | 1,101 | 1,442 | 28% | 0.902 | 0.967 | 0.948 | 0.915 | 0.908 |
| **LICKOB**<br>**JECT** | 460 | 629 | 1.8% | 0.410 | 0.990 | 0.98 | 0.439 | 0.4<br>24 |
| **LICKSE**<br>**LF** | 257 | 398 | 3.4% | 0.772 | 0.990 | 0.982 | 0.728 | 0.7<br>49 |
| **PETTIN**<br>**G** | 204 | 307 | 0.96% | 0.305 | 0.991 | 0.984 | 0.237 | 0.2<br>67 |
| **RUBBIN**<br>**G** | 158 | 235 | 0.73% | 0.729 | 0.996 | 0.994 | 0.584 | 0.6<br>48 |
| **SCRAT**<br>**CH** | 158 | 303 | 0.60% | 0.870 | 0.997 | 0.997 | 0.676 | 0.7<br>61 |
| **SHAKE** | 251 | 435 | 0.13% | 0.916 | 1.00 | 1.00 | 0.795 | 0.8<br>51 |
| **SNIFF** | 946 | 1,74<br>7 | 5.3% | 0.610 | 0.968 | 0.949 | 0.517 | 0.5<br>59 |
| **NONE** | 2,051 | 4,63<br>6 | 55% | 0.892 | 0.898 | 0.895 | 0.914 | 0.9<br>03 |
| **Posture** |  |  |  |  |  |  |  |  |
| **LIE**<br>**DOWN** | 674 | 1,22<br>3 | 22% | 0.826 | 0.913 | 0.894 | 0.724 | 0.7<br>72 |
| **SIT** | 726 | 1,27<br>5 | 9.9% | 0.409 | 0.915 | 0.865 | 0.347 | 0.3<br>75 |
| **STAND** | 2,028 | 4,24<br>1 | 58% | 0.793 | 0.900 | 0.838 | 0.916 | 0.8<br>50 |
| **VIGOROUS** | 289 | 468 | 2.9% | 0.764 | 0.985 | 0.978 | 0.605 | 0.675 |
| **WALK** | 599 | 905 | 7.7% | 0.903 | 0.969 | 0.964 | 0.706 | 0.793 |

**Table 7.**
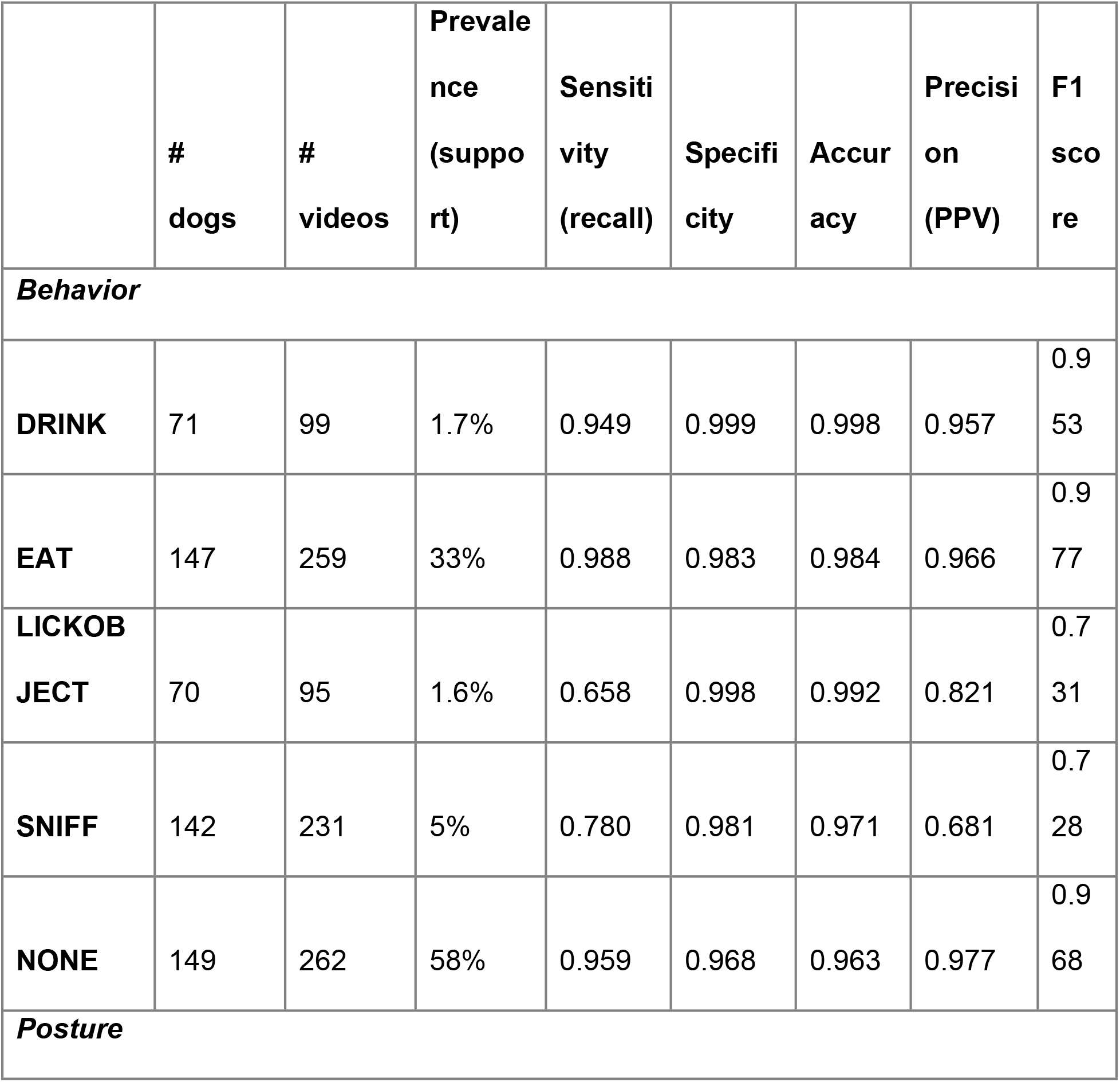
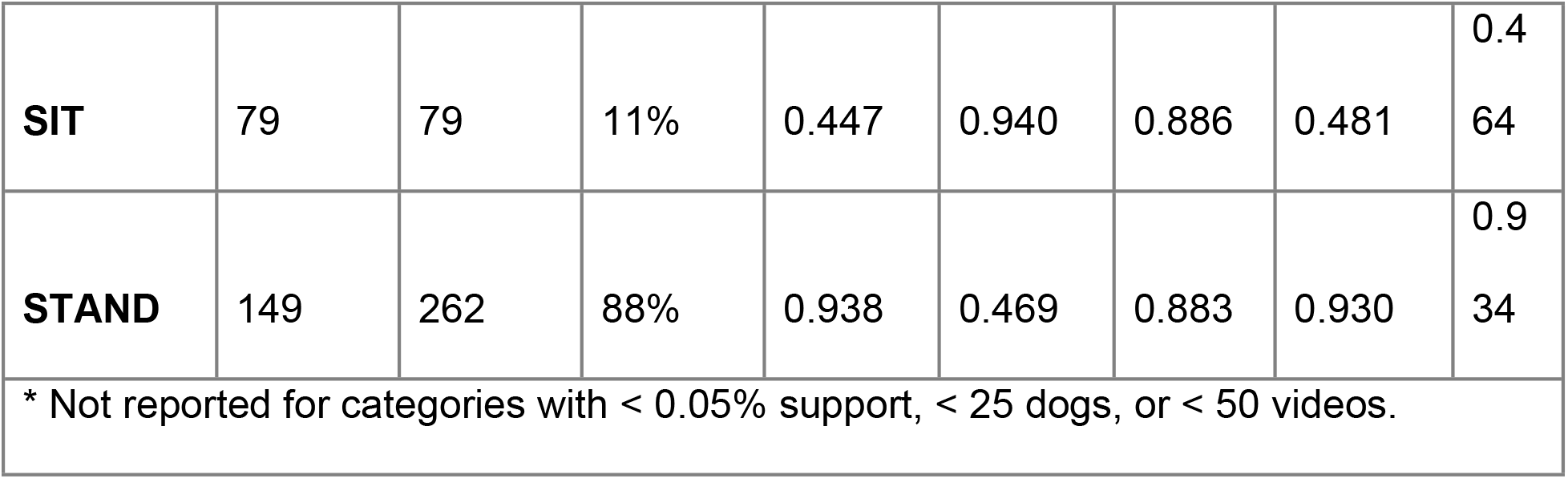
Classification metrics for the *eat/drink* dataset.*

The “behaviors” confusion matrix for the *crowd* dataset is shown in Fig 6 in nonnormalized and normalized forms. The non-normalized confusion matrix gives raw tallies (that is, the total number of one-third second time points) of predicted and labeled classes, and the normalized confusion matrix gives the percentage of each actual label classified by the algorithms as a given predicted label (so that the percentages in each row sum to 100%). The non-normalized matrix is dominated by correctly predicted NONE and EAT samples, due to their high prevalence and effective classification in this dataset. The normalized matrix suggests the reliable classification of DRINK, EAT, NONE, and SHAKE. The LICKSELF and SCRATCH classes are of moderate reliability, and the LICKOBJECT, PETTING, RUBBING, and SNIFF classes exhibit some systematic misclassification and are of lesser reliability.

**Fig 6.**
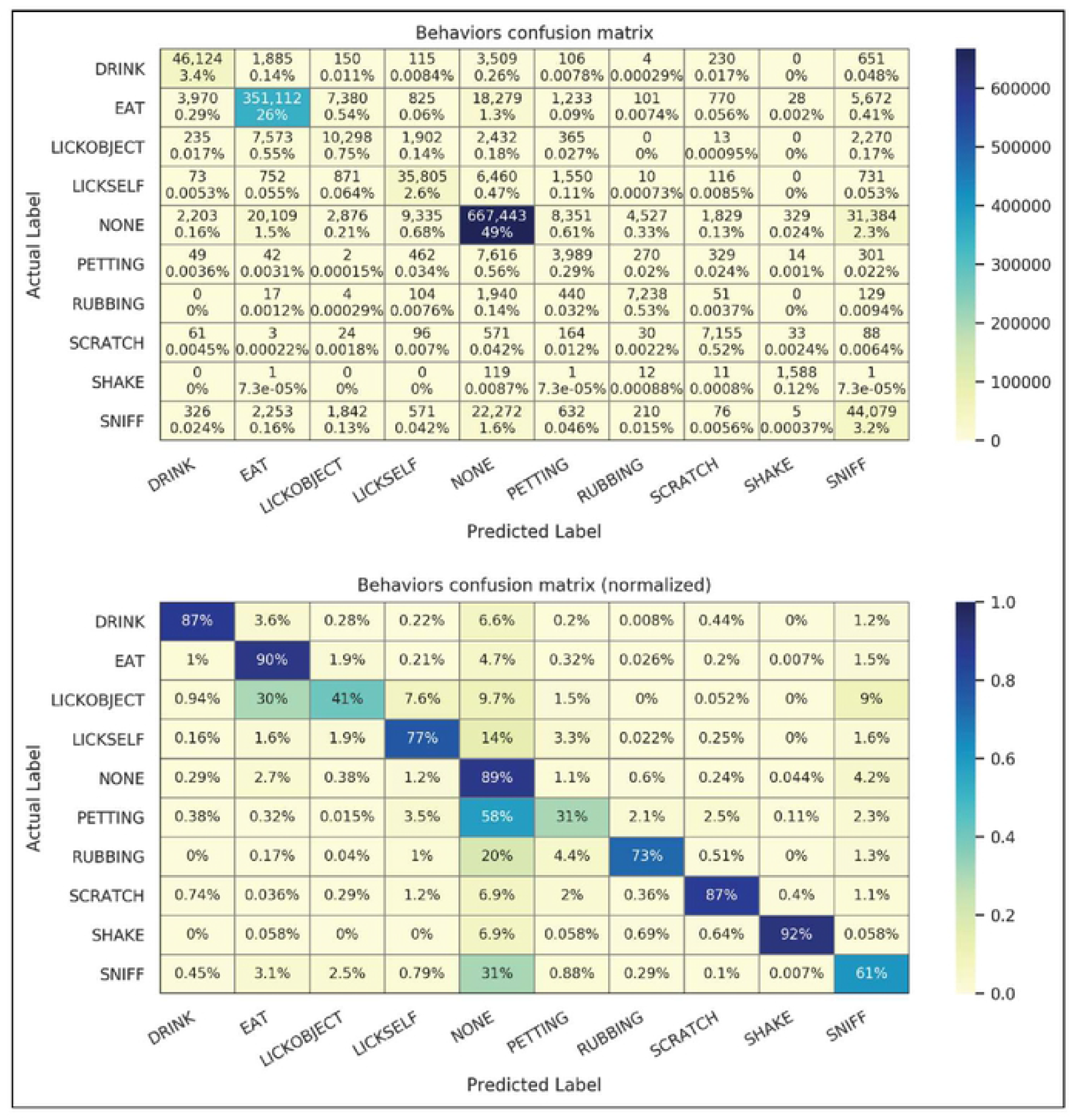
Confusion matrices for *crowd* dataset behaviors. Timepoint values in raw counts (top) and normalized by timepoint labels (bottom) are shown.

### Effect of device position on performance

The system’s classification performance, as measured by F1 score, shows no significant dependence on device position (Fig 7). This invariance is a key property that enables real-world performance to approach that of controlled studies.

**Fig 7.**
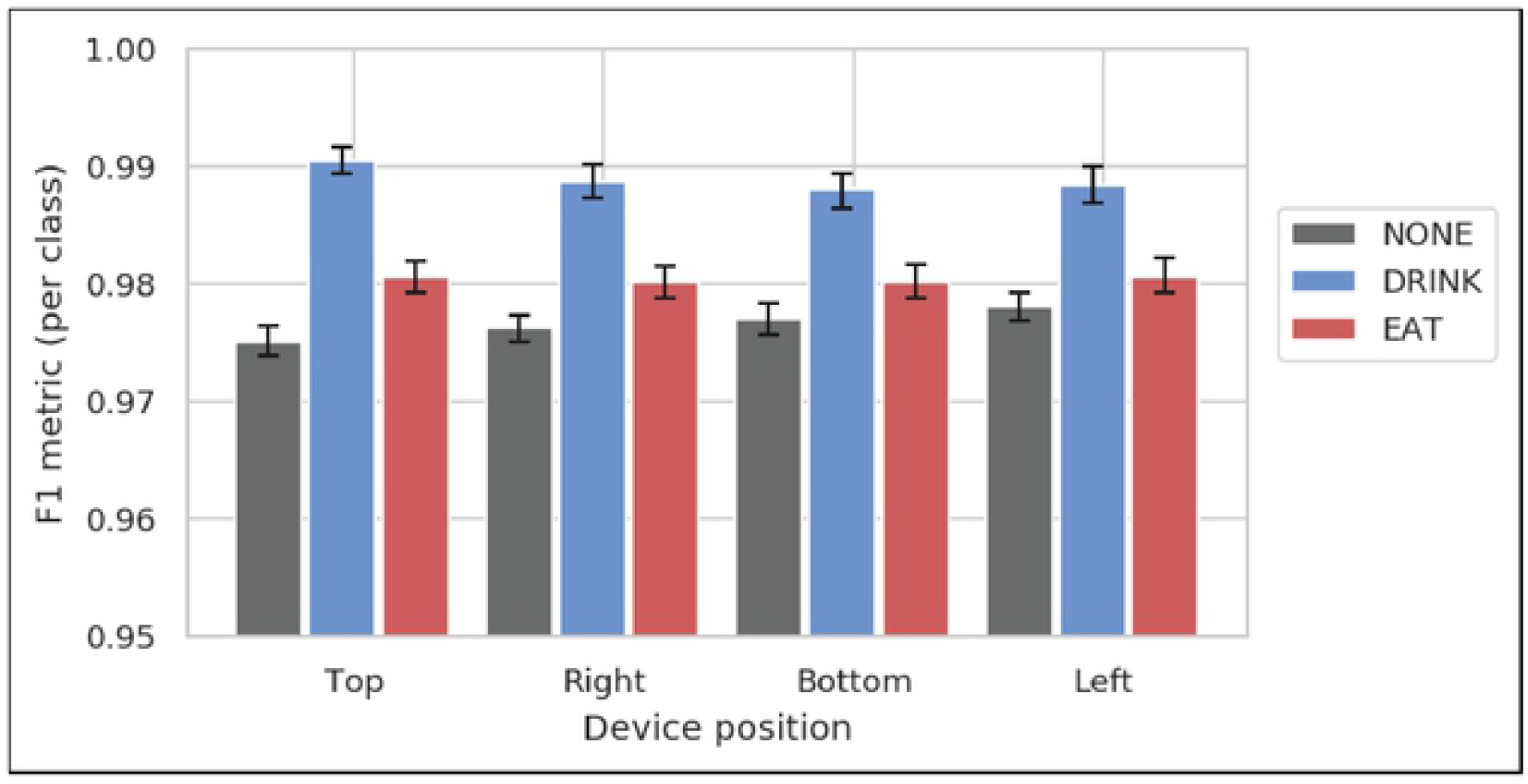
Classification performance as measured by F1 score. F1 scores measuring test accuracy are broken out by device position, for n=48 videos from the *eat/drink* dataset where the dog’s collar had exactly four attached devices with known orientation. Error bars are 95% confidence intervals on the mean, as determined by bootstrapping. The classification accuracy per class was similar between the four positions, indicating that system accuracy is not substantially affected by collar rotation.

### User validation

Participants responded far better than expected to user validation efforts. Users opened emails, clicked through to the web form, and submitted validation results for 55% of the EAT validation emails and 42% of the DRINK validation emails.

Responses are summarized in Table 8. As described above, we excluded any responses that arrived more than 60 minutes after an event’s end, as well as any “Not Sure” responses. The positive (“Yes”) validation rate was approximately 95% for both event types. As expected, the rate of users responding “Not Sure” was far greater for DRINK (12%) than for EAT (2%). Participant comments confirmed our expectation that users were less aware of DRINK behavior than of EAT behavior. This lack of awareness likely also contributed to the lower DRINK response rate.

**Table 8.**
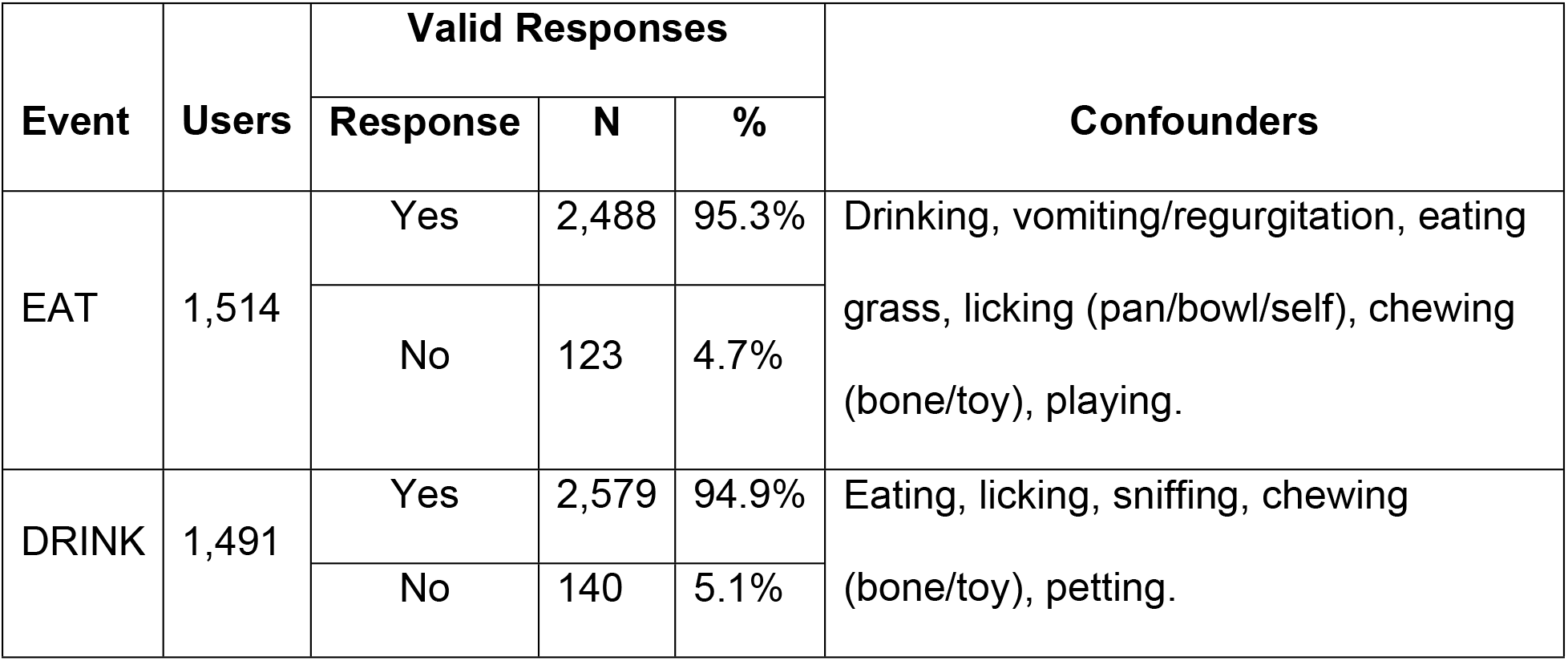
Summary of user EAT and DRINK validation responses.

As the production system generates candidate EAT and DRINK events, it calculates a confidence score (the mean algorithm confidence over the event’s duration) that varies between 0 and 1.0, and drops any events with a score below a threshold of 0.3. Fig 8 shows how the percentage of “Yes” responses (the true positive rate) varied with this confidence score. For EAT events, the rate grew from 83% for the lowest-confidence bin (0.3-0.4) to 100% (201 out of 201) for the highest-confidence bin (0.9-1.0). Since users do not see the confidence score, this trend suggests that the EAT validation data are relatively reliable. The DRINK data show a less convincing trend, which is consistent with users’ lower awareness of DRINK events.

**Fig 8.**
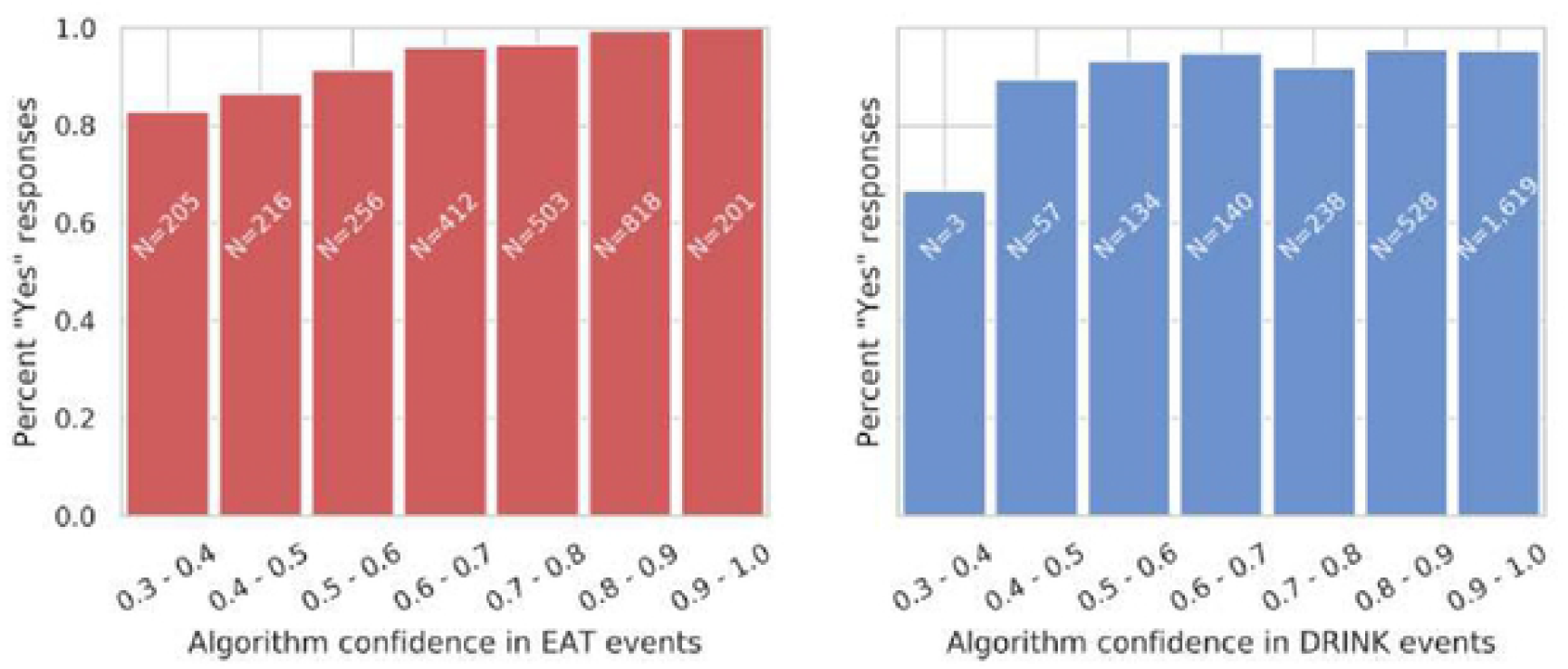
User responses to EAT and DRINK event validation requests. Responses to EAT events (left) and DRINK events (right) are grouped by the algorithm’s confidence in each event. Validation requests were sent automatically by email within 15 minutes of an event, and responses were allowed within 1 hour of an event. Confidence estimates varied from 0.3 to 1.0 (events with lower confidence are dropped by the algorithm). Bars are annotated with the total number of responses in each bin.

It is unfortunate that, of the behavior classes measured in this work, only EAT is likely to exhibit the level of user awareness required for validation using this method.

## Discussion

### Comparison with Previous Work

We compare our dataset and results with several previous works in Table 9, and we tabulate several important qualitative differences between the datasets in Table 10.

In comparing these results, it is important to account for:

1. **Class distribution**. Each dataset exhibits a different distribution of behaviors. In general, classifiers exhibit better F1 scores for common behaviors than for rare behaviors. The classifier sensitivity and specificity are relatively insensitive to this distribution, so we recommend using these metrics for comparing performance across different datasets.
2. **Dataset collection methods**. Classifiers are more accurate when applied to high-quality datasets collected under controlled conditions. Accuracy can drop substantially in naturalistic versus laboratory settings [26,27]. Classifiers benefit from consistent device position, device attachment, and collar tightness, and they also benefit when the labeled behaviors as well as the collection environment are consistent and well-defined.

**Table 9.**
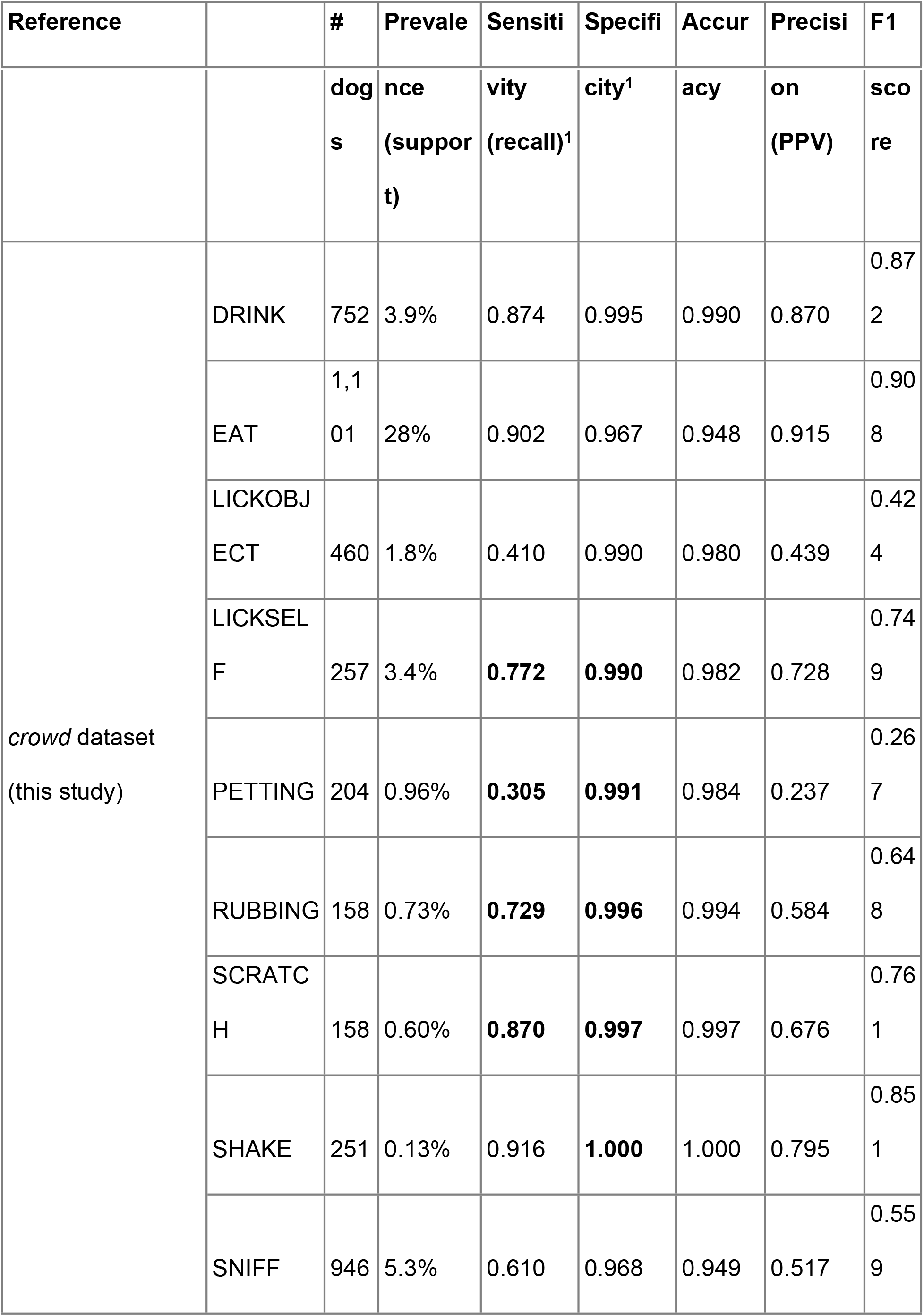
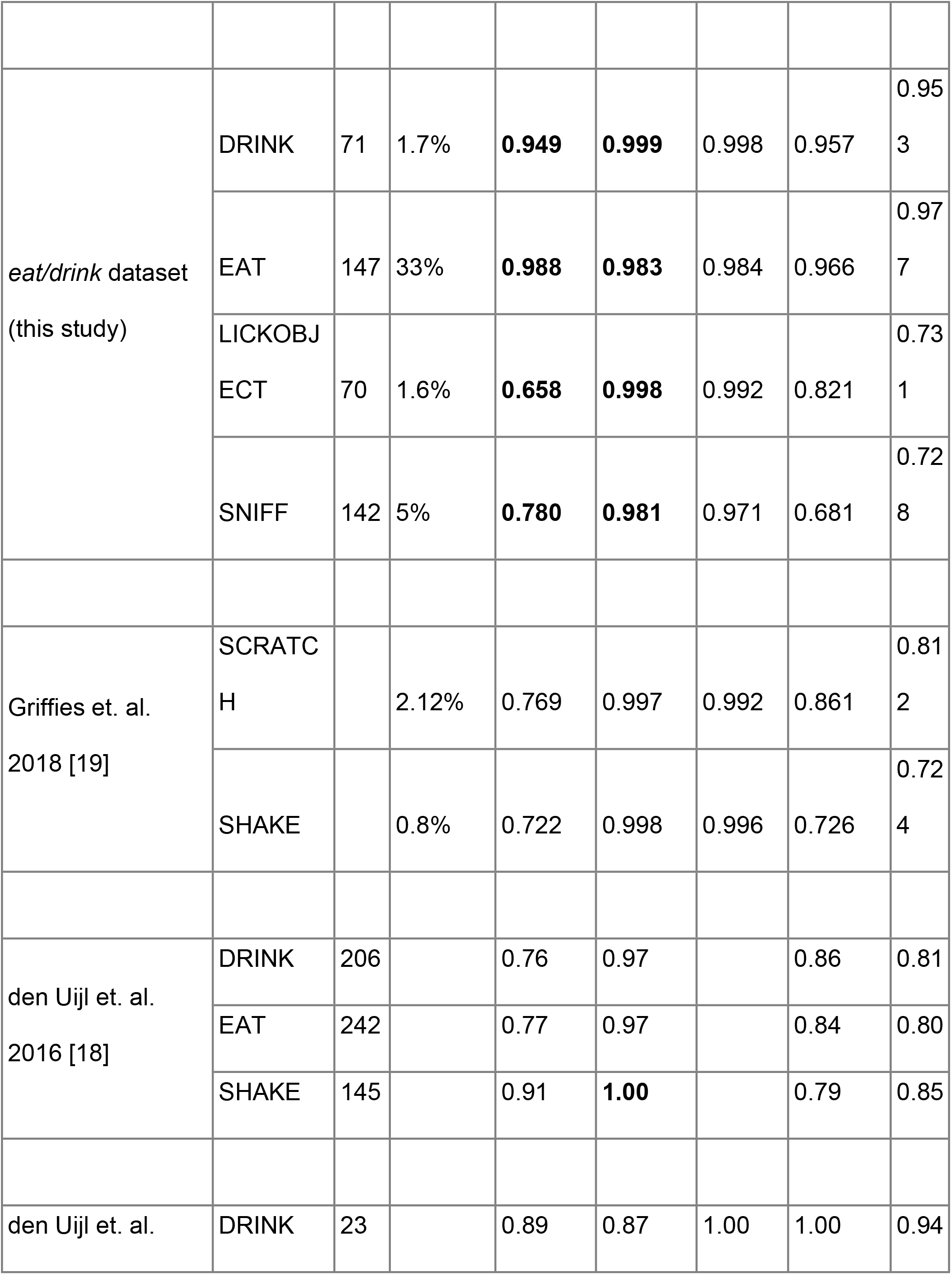
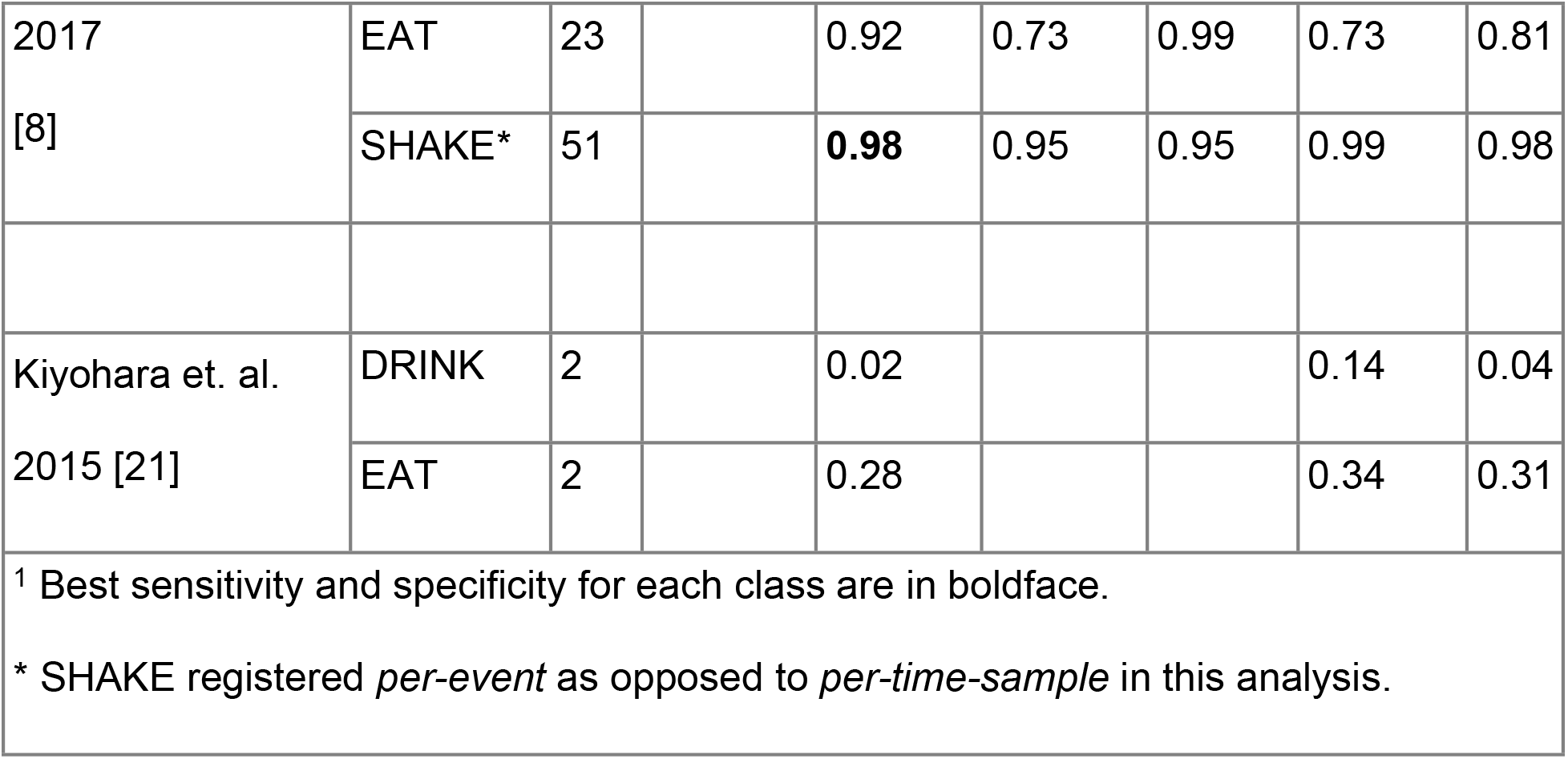
Comparison of this work to other published results.

**Table 10.** Dataset collection methods in similar published studies.

| Reference* | Device position | Collar fit | Environment | Behaviors |
| --- | --- | --- | --- | --- |
| *Crowd* dataset<br>(this study) | Uncontrolled | Uncontrolled | Uncontrolled<br>(highly varied) | Naturalistic |
| *Eat/drink* dataset<br>(this study) | Controlled | Controlled | Controlled<br>(lab/kennel) | Semi-controlled |
| Griffies et. al.<br>2018 [19] | Controlled | Controlled | Controlled<br>(humane society) | Naturalistic |
| den Uijl et. al.<br>2017 [8] | Controlled | Controlled | Controlled<br>(track/field and<br>indoors) | Semi-controlled |
| * Relevant data collection details not available for den Uijl et. al. 2016 [18] or |  |  |  |  |

Previous works have used relatively controlled and high-quality datasets, similar to the *eat/drink* dataset in this work [8,18,19,21]. The classification performance of the classifier presented here on the EAT and DRINK classes in the *eat/drink* dataset advances the sensitivity, specificity, and F1 score for these classes. Sensitivity and specificity are independent of class prevalence. The balance between sensitivity and specific is a design choice, so we have calibrated our algorithms to favor specificity in order to minimize false positives.

The classifiers’ performance on SCRATCH in the challenging *crowd* dataset also advances the state of the art. Comparable detection of LICKOBJECT, LICKSELF, PETTING, RUBBING, and SNIFF has not been previously demonstrated to our knowledge. We note that SCRATCH, LICKSELF, and RUBBING behaviors are highly relevant to dermatological health applications [19], and that PETTING is an important confounder that can be easily misclassified as SCRATCH or LICKSELF in classifiers that are not exposed to this behavior. We have found the classifiers’ detection of SHAKE to be highly accurate (though susceptible to temporal misalignment between device and video data, due to the short event lengths). It is difficult to compare the *per-time-sample* SHAKE classification metrics here to published *per-event* metrics due to differing methodologies [8,18].

### Challenges

The system excels at certain clearly defined and easily recognizable activities, especially those repeating and universal movement patterns such as drinking (lapping), walking, running, shaking, and most eating behaviors. It also performs well on well-defined instances of scratching and self-licking.

Device positioning and collar tightness do not appear to have a strong effect on system accuracy, meaning that accurate behavior metrics can be acquired via normal activity monitor usage. An important feature of the devices described in this study is their insensitivity (invariance) to collar orientation or position (Fig 7). In real-world settings, and especially with lightweight devices such as the Whistle FIT, the device can be, and often is, rotated away from the conventional ventral (bottom) position at the lowest point of the collar.

The system appears to use the angle of a dog’s neck (that is, whether the dog is looking up or down) as an important behavioral clue. Consequently, activities such as eating or drinking appear to be less accurate when raised dog bowls are used, and activities such as sniffing and scratching, and self-licking can go undetected if performed in unusual positions. Slow-feed food bowls, collars attached to taut leashes, and loose collars with other heavy attachments can also cause misclassifications, but are often classified correctly nonetheless.

Other sets of activities simply present very similar accelerometer data, such as eating wet food, which can be confounded with drinking; or being pet by a human or riding in a moving vehicle, which can be confounded with scratching or self-licking; or even vigorous playing and ‘tug-of-war’, which can be confounded with shaking and other activities. These misclassifications become less common as the models improve, but in some cases confusion may be unavoidable. Some other activities are simply rare or unusual, for instance, drinking from a stream, drinking from a water bottle, or licking food off of a raised plate.

A different type of problem relates to activities that are ambiguous even to human labelers, such as the distinction between eating a small part of a meal versus eating a large treat. Similarly, when a dog repeatedly starts and stops an activity, it is often a matter of the labeler’s judgment whether to use a single long label or multiple short labels. Both of these types of labeling ambiguity can be very deleterious to certain classification metrics, even though it is questionable whether the system’s usefulness or real-world accuracy is affected.

## Conclusion

We advanced the sensitivity and specificity for detecting drinking and eating behavior, and we demonstrate detection of licking, petting, rubbing, scratching, and sniffing, which to our knowledge have not been reported in a comparable manner. We demonstrated that system performance is not sensitive to collar position. In production, users reported high rates of true positives, consistent with the metrics measured via cross-validation on the *crowd* training database. The systems described in this work can further improve via the incorporation of additional training data and through the improvement of the underlying algorithms.

## Acknowledgments

We are grateful to Leonid Sudakov and Jeannine Taaffe for their support and vision in enabling the Pet Insight Project; WALTHAM Petcare Science Institute for contributing extensive training data and for numerous helpful discussions; to the many participants in the Pet Insight Project; and to the Whistle, Kinship, and Mars Petcare organizations for supporting the development and publication of this work.

## Author contributions

R.C: Conceptualization, Data Curation, Formal Analysis, Investigation, Methodology, Software, Validation, Visualization, Writing – Original Draft Preparation

N.Y: Conceptualization, Data Curation, Formal Analysis, Investigation, Methodology, Software, Validation, Writing – Review & Editing

A.C Conceptualization, Investigation--in-clinic crowd sourcing, Methodology, Supervision, Writing – Review & Editing

C. J: Data Curation, Investigation, Methodology, Software, Writing – Review & Editing

D. A: Conceptualization, Data Curation, Investigation, Validation, Project Administration L.P: Software, Writing – Review & Editing

S. B: Conceptualization support -- eating and drinking study, Investigation--performed the experimental work at the WALTHAM Petcare Science Institute, Writing – Review & Editing

G.W: Conceptualization, Funding Acquisition, Methodology, Project Administration

K.L: Conceptualization, Funding Acquisition, Methodology, Resources

S.L: Conceptualization, Writing – Review & Editing

## References

1. Pewek L, Ellis DA, Andrews S, Joinson A. Wearables: Promises and Barriers. 2016; PLoS Med. 13: doi:10.1371/journal.pmed.1001953.

2. Vogenberg FR, Santilli J. Healthcare trends for 2018. 2018; Am Health Drug Benefits. 11: 48–54.

3. Tison GH, Sanchez JM, Ballinger B, Singh A, Olgin JE, Pletcher MJ, et al. Passive detection of atrial fibrillation using a commercially available smartwatch. 2018. JAMA Cardiol. 2018;3: 409–416. doi:10.1001/jamacardio.2018.0136.

4. Pramanik PKD, Upadhyaya BK, Pal S, Pal T. Chapter 1 – Internet of things, smart sensors, and pervasive systems: Enabling connected and pervasive healthcare. In: Dey N, Ashour AS, Bhatt C, James Fong S, editors. Healthcare Data Analytics and Management. London: Academic Press; 2019. pp. 1–58. doi:10.1016/B978-0-12-815368-0.00001-4.

5. Watson K, Wells J, Sharma M, Robertson S, Dascanio J, Johnson JW, et al. A survey of knowledge and use of telehealth among veterinarians. 2019; BMC Veterinary Research. 15: 474. doi:10.1186/s12917-019-2219-8.

6. Pacis DMM, Subido EDC, Bugtai NT. Trends in telemedicine utilizing artificial intelligence. AIP Conference Proceedings. 2018; 1933: 040009. doi:10.1063/1.5023979.

7. Kour H, Patison KP, Corbet NJ, Swain DL. Validation of accelerometer use to measure suckling behaviour in Northern Australian beef calves. 2018; Appl Anim Behav Sc. 202: 1–6. doi:10.1016/j.applanim.2018.01.012.

8. den Uijl I, Gómez Álvarez CB, Bartram D, Dror Y, Holland R, Cook A. External validation of a collar-mounted triaxial accelerometer for second-by-second monitoring of eight behavioural states in dogs. Wade C, editor. PLOS ONE. 2017; 12: e0188481. doi:10.1371/journal.pone.0188481.

9. Belda B, Enomoto M, Case BC, Lascelles BDX. Initial evaluation of PetPace activity monitor. 2018; Vet J. 237: 63–68. doi:10.1016/j.tvjl.2018.05.011.

10. Weiss GM, Nathan A, Kropp JB, Lockhart JW. WagTag: a dog collar accessory for monitoring canine activity levels. Proceedings of the 2013 ACM conference on Pervasive and ubiquitous computing adjunct publication. UbiComp ‘13 Adjunct. Zurich, Switzerland: ACM Press; 2013. pp. 405–414. doi:10.1145/2494091.2495972.

11. Mejia S, Duerr FM, Salman M. Comparison of activity levels derived from two accelerometers in dogs with osteoarthritis: Implications for clinical trials. 2019; Vet J. 252: 105355. doi:10.1016/j.tvjl.2019.105355.

12. Westgarth C, Ladha C. Evaluation of an open source method for calculating physical activity in dogs from harness and collar based sensors. 2017; BMC Vet Res. 13: 322. doi:10.1186/s12917-017-1228-8.

13. Hansen BD, Lascelles BDX, Keene BW, Adams AK, Thomson AE. Evaluation of an accelerometer for at-home monitoring of spontaneous activity in dogs. 2007; Am J Vet Res. 68: 468–475. doi:10.2460/ajvr.68.5.468.

14. Hoffman CL, Ladha C, Wilcox S. An actigraphy-based comparison of shelter dog and owned dog activity patterns. 2019; J Vet Behavior. 34: 30–36. doi:10.1016/j.jveb.2019.08.001.

15. Kumpulainen P, Valldeoriola A, Somppi S, Törnqvist H, Väätäjä H, Majaranta P, et al. Dog activity classification with movement sensor placed on the collar. Proceedings of the Fifth International Conference on Animal-Computer Interaction. Atlanta, Georgia, USA: Association for Computing Machinery; 2018. pp. 1–6. doi:10.1145/3295598.3295602.

16. Brugarolas R, Loftin RT, Yang P, Roberts DL, Sherman B, Bozkurt A. Behavior recognition based on machine learning algorithms for a wireless canine machine interface. 2013 IEEE International Conference on Body Sensor Networks. 2013. pp. 1–5. doi: 10.1109/BSN.2013.6575505.

17. Petrus S, Roux L. Real-time behaviour classification techniques in low-power animal borne sensor applications. Thesis, Stellenbosch: Stellenbosch University. 2019. Available: https://scholar.sun.ac.za:443/handle/10019.1/105744.

18. den Uijl I, Gomez-Alvarez C, Dror Y, Manning N, Bartram D, Cook A. Validation of a collar-mounted accelerometer that identifies eight canine behavioural states, including those with dermatologic significance. Proceedings of British Beterinary Dermatology Study Group. Weybridge UK; 2016. pp. 81–84.

19. Griffies JD, Zutty J, Sarzen M, Soorholtz S. Wearable sensor shown to specifically quantify pruritic behaviors in dogs. 2018; BMC Vet Res. 14: 124. doi:10.1186/s12917-018-1428-x.

20. Dog’s life. Proceedings of the 2013 ACM international joint conference on Pervasive and ubiquitous computing. [cited 10 Jan 2020]. Available: https://dl.acm.org/doi/abs/10.1145/2493432.2493519.

21. Kiyohara T, Orihara R, Sei Y, Tahara Y, Ohsuga A. Activity recognition for dogs based on time-series data analysis. In: Duval B, van den Herik J, Loiseau S, Filipe J, editors. Agents and Artificial Intelligence. Cham: Springer International Publishing; 2015. pp. 163–184. doi:10.1007/978-3-319-27947-3_9.

22. Nuttall T, McEwan N. Objective measurement of pruritus in dogs: a preliminary study using activity monitors. 2006; Vet Dermatol. 17: 348–351. doi:10.1111/j.1365-3164.2006.00537.x.

23. Plant JD. Correlation of observed nocturnal pruritus and actigraphy in dogs. 2008; Vet Rec. 162: 624–625. doi:10.1136/vr.162.19.624.

24. Morrison R, Penpraze V, Beber A, Reilly JJ, Yam PS. Associations between obesity and physical activity in dogs: a preliminary investigation. 2013; J Small Anim Pract. 54: 570–574. doi:10.1111/jsap.12142.

25. Helm J, McBrearty A, Fontaine S, Morrison R, Yam P. Use of accelerometry to investigate physical activity in dogs receiving chemotherapy. 2016; J Small Anim Pract. 57: 600–609. doi:10.1111/jsap.12587.

26. Twomey N, Diethe T, Fafoutis X, Elsts A, McConville R, Flach P, et al. A comprehensive study of activity recognition using accelerometers. 2018; Informatics. 5: 27. doi:10.3390/informatics5020027.

27. Foerster F, Smeja M, Fahrenberg J. Detection of posture and motion by accelerometry: a validation study in ambulatory monitoring. 1999; Computers Human Behavior. 15: 571–583. doi:10.1016/S0747-5632(99)00037-0.

28. Olsen AM, Evans RB, Duerr FM. Evaluation of accelerometer inter-device variability and collar placement in dogs. 2016; Vet Evidence. doi:10.18849/ve.v1i2.40.

29. Martin KW, Olsen AM, Duncan CG, Duerr FM. The method of attachment influences accelerometer-based activity data in dogs. 2017; BMC Vet Res. 13: 48. doi:10.1186/s12917-017-0971-1.

30. Aich S, Chakraborty S, Sim J-S, Jang D-J, Kim H-C. The design of an automated system for the analysis of the activity and emotional patterns of dogs with wearable sensors using machine learning. 2019; Applied Sciences. 9: 4938. doi:10.3390/app9224938.

31. Pet Insight Project. In: Pet Insight Project [Internet]. [cited 16 Dec 2019]. Available: https://www.petinsight.co.

32. Chambers RD, Yoder NC. FilterNet: A many-to-many deep learning architecture for time series classification. 2020; Sensors. 20: 2498. doi:10.3390/s20092498.

33. Friard O, Gamba M. BORIS: a free, versatile open-source event-logging software for video/audio coding and live observations. 2016; Methods Ecol Evolution. 7: 1325–1330. doi:10.1111/2041-210X.12584.

34. Hammerla NY, Plötz T. Let’s (not) stick together: pairwise similarity biases crossvalidation in activity recognition. Proceedings of the 2015 ACM International Joint Conference on Pervasive and Ubiquitous Computing. Osaka, Japan: Association for Computing Machinery; 2015. pp. 1041–1051. doi:10.1145/2750858.2807551.

35. Gerencsér L, Vásárhelyi G, Nagy M, Vicsek T, Miklósi A. Identification of behaviour in freely moving dogs (Canis familiaris) using inertial sensors. 2013; PLOS ONE. 8: e77814. doi:10.1371/journal.pone.0077814

36. Paszke A, Gross S, Massa F, Lerer A, Bradbury J, Chanan G, et al. PyTorch: An imperative style, high-performance deep learning library. In: Wallach H, Larochelle H, Beygelzimer A, Alché-Buc F, Fox E, Garnett R, editors. Advances in Neural Information Processing Systems 32. Curran Associates, Inc.; 2019. pp. 8024–8035. Available: http://papers.nips.cc/paper/9015-pytorch-an-imperative-style-high-performance-deep-learning-library.pdf.

37. Amazon EC2-P2 Instances. In: Amazon Web Services, Inc. [Internet]. [cited 6 Dec 2019]. Available: https://aws.amazon.com/ec2/instance-types/p2/.

38. Haghighi S, Jasemi M, Hessabi S, Zolanvari A. PyCM: Multiclass confusion matrix library in Python. 2018; JOSS. 3: 729. doi:10.21105/joss.00729.

39. Steyerberg E. Clinical Prediction Models: A Practical Approach to Development, Validation, and Updating. New York: Springer-Verlag; 2009. doi:10.1007/978-0-387-77244-8.

